# From *Nicotiana* to apple: effector screening reveals new avenues for durable scab resistance

**DOI:** 10.64898/2025.12.25.696553

**Authors:** Silvia de la Rosa, Mariana Tarallo, Joanna Tannous, Mélanie Sannier, Yi-Hsuan Tu, Maël Baudin, Joanna K. Bowen, Ashleigh M. Mosen, Kim M. Plummer, Rosie E. Bradshaw, Bruno Le Cam, Carl H. Mesarich

**Author notes:** Corresponding authors: Carl H. Mesarich and Bruno Le Cam. Biosciences Division, Oak Ridge National Laboratory, Oak Ridge, Tennessee, United States of America (Joanna Tannous).

## Abstract

Scab or black spot disease, caused by the fungus *Venturia inaequalis*, is an ongoing threat to commercial apple production. Current control methods involve extensive fungicide use and the deployment of disease-resistant apple cultivars. However, fungicide-resistant strains of *V. inaequalis* are becoming more prevalent, as are strains that can overcome one or more qualitative disease resistance genes in apple. To increase the durability of disease resistance, and thus decrease our reliance on fungicides, one promising approach could involve stacking endogenous and exogenous resistance genes in apple cultivars using genetic modification. As a starting point for the identification of exogenous resistance genes that are effective against *V. inaequalis*, 137 candidate effector (CE) proteins from this fungus, fused to a signal peptide for extracellular targeting to the apoplast, were screened for recognition by extracellular leucine-rich repeat (LRR)–receptor-like protein and LRR–receptor-like kinase immune receptors in the model angiosperm species *Nicotiana benthamiana* and *Nicotiana tabacum* using *Agrobacterium tumefaciens*-mediated transient transformation assays. Here, a cell death response in wild-type plants, together with a loss of this response in plants lacking the extracellular immune co-receptor *Nb*SOBIR1 or *Nb*BAK1, was used as an indicator of recognition. In total, six CE proteins triggered cell death in one or both *Nicotiana* species, but only one, a homolog of the *Vm*E02 effector protein from the apple pathogen *Valsa mali*, did so in an *Nb*SOBIR1- and *Nb*BAK1-dependent manner. The five remaining CE proteins are homologs of other known cell death elicitors from filamentous plant pathogens for which there is evidence that they trigger non-canonical extracellular immunity in plants. One of these is an Alt a 1-like protein that also triggered cell death in apple. Collectively, these findings provide a foundation for the use of a combined set of exogenous and endogenous resistance genes in apple to provide durable protection against scab disease.

## Introduction

Scab or black spot disease of apple (*Malus* × *domestica*), caused by the biotrophic fungus *Venturia inaequalis*, is an ongoing threat to commercial apple production worldwide [1]. Inadequate management of this disease can result in economic losses ranging from 70 to 100% [2]. These losses are primarily due to fruit infections, which lead to premature fruit drop and blemished produce that is no longer marketable, as well as leaf infections, which decrease the growth and yield of apple trees through premature leaf drop and reduced photosynthetic capacity [3]. To mitigate these losses, orchardists typically rely on 12 to 25 fungicide applications per season [2]. However, while fungicides are typically effective at controlling scab disease, fungicide-resistant strains of *V. inaequalis* are becoming increasingly prevalent [4]. This issue, together with the environmental harm associated with fungicide use, and the growing concerns over fungicide residues on fruit, indicate that reliance on this approach is not sustainable in the long term and calls for the development of novel control approaches.

As an alternative to fungicides, scab disease can be controlled through the cultivation of resistant apple varieties carrying *Rvi* (*Resistance to Venturia inaequalis*) genes, which confer major, qualitative resistance [5]. To date, 19 *Rvi* genes have been genetically identified in apple [5, 6] and, of these, three have so far been cloned to determine the type of immune receptors they encode. These are *Rvi15*, recently shown to be the same gene as *Rvi4* [7], which encodes an intracellular Toll interleukin-1 receptor-like nucleotide-binding site–leucine-rich repeat (TIR-NBS-LRR) protein [8], *Rvi6*, which encodes an extracellular membrane-bound LRR–receptor-like protein (LRR-RLP) [9], and *Rvi12*, which encodes an extracellular membrane-bound LRR–receptor-like kinase (LRR-RLK) protein [10]. To activate the defence responses that are necessary for scab resistance, Rvi immune receptors must directly or indirectly recognize specific effectors of *V. inaequalis*. Since this recognition often renders *V. inaequalis* unable to cause disease, the recognized effectors are termed avirulence effectors and are encoded by *AvrRvi* (*Avirulence to Rvi*) genes [5]. Apart from *AvrRvi6*, which corresponds to *Rvi6* [11], the *AvrRvi* genes of *V. inaequalis* corresponding to the *Rvi* genes of apple have not yet been cloned or published.

Typically, the defence responses that develop following effector recognition involve a localized form of cell death, called the hypersensitive response (HR), which is based on chlorosis or necrosis [5]. The type and strength of HR vary according to the specific Rvi immune receptor involved [5]. Additionally, many minor resistance genes associated with quantitative trait loci (QTLs) have been genetically identified in apple, and these confer a low level of protection against scab disease [6]. The mechanisms by which these QTLs provide protection remain unknown. However, this could conceivably also involve effector recognition [12], or the recognition of other molecular patterns associated with host invasion (i.e., conserved pathogen-associated molecular patterns (PAMPs) or endogenous damage-associated molecular patterns (DAMPs)), by extracellular LRR-RLP or LRR-RLK immune receptors [13].

In the absence of corresponding Rvi immune receptors, the effectors of *V. inaequalis* function as virulence factors in promoting host colonization and disease. Based on what is known from other plant-pathogenic fungi [14, 15], most of these effectors are expected to be secreted proteins. The exact number of secreted proteins produced by *V. inaequalis* is predicted to range from 647 to 1,955, varying with the strain studied, the bioinformatic resource used (i.e., genome or transcriptome sequence), and the computational pipeline employed [16–20]. Nevertheless, relatively little research has been conducted on candidate effector (CE) proteins of *V. inaequalis* [11, 19, 21–25]. Consequently, their specific roles in virulence remain unknown. What is known, however, is that most CE proteins of *V. inaequalis* are less than 300 amino acid residues in length and belong to expanded families of two or more members [16, 19]. Furthermore, the genes encoding these CE proteins are upregulated as part of temporal waves of expression during infection [16, 19]. For example, AvrRvi6 belongs to an expanded family of approximately 75 members with structural similarity to the MAX (Magnaporthe AVRs and ToxB-like) effectors of the rice pathogen *Magnaporthe oryzae* and is encoded by a gene that peaks in expression at 3–5 days post-inoculation of susceptible apple [11, 19].

Unfortunately, as with fungicide-based control, single *Rvi* genes in apple can be overcome by *V. inaequalis*. This breakdown is reflected in data from the VINQUEST project (https://www.vinquest.ch), which monitors the geographical distribution of *V. inaequalis* isolates across 15 differential apple hosts carrying single *Rvi* genes. Indeed, based on field observations from 24 project partners across 14 countries in the 10 years following the initiation of VINQUEST (i.e., 2009–2018), virulent isolates of *V*. *inaequalis* were detected against most *Rvi* genes [26]. Virulences towards *Rvi1*, *Rvi3*, and *Rvi8* are the most prevalent, with the high level of *Rvi1* breakdown likely reflecting the extensive deployment of this gene in apple cultivars [27]. *V. inaequalis* is predicted to overcome these *Rvi* genes primarily through deletion or mutation of the corresponding *AvrRvi* genes. Indeed, mutations leading to amino acid substitutions in AvrRvi6 enable the pathogen to circumvent *Rvi6*-mediated resistance [11]. Unexpectedly, molecular dating of these events within virulent European populations revealed that they predate apple domestication, with wild endemic *Malus* species acting as long-term reservoirs of virulence. The extensive diversity of wild *Malus* species across the Northern Hemisphere raises the possibility that they already harbour virulent populations of *V. inaequalis* capable of overcoming other *Rvi* genes, potentially compromising breeding efforts. Consequently, diversifying the host origin of resistance genes, particularly by incorporating those that have never co-evolved with *V. inaequalis*, is of primary importance for achieving durable resistance.

*Agrobacterium tumefaciens*-mediated transient transformation assays (ATTAs) have successfully been employed to demonstrate that a portion of the effectors within the secretomes of diverse plant-pathogenic fungi trigger a cell death response when expressed in *Nicotiana* species [28–33]. In many cases, this response is dependent on the possession of a signal peptide for extracellular targeting of the CE proteins to the apoplast, as well as one or both of the plant co-receptors, BAK1 and SOBIR1, which mediate downstream immune responses following the recognition of extracellular CE proteins, PAMPs and DAMPs by LRR-RLPs (which are dependent on both BAK1 and SOBIR1) and LRR-RLKs (which are dependent on only BAK1) [34–39]. For example, 14 of 63 CE proteins tested from the wheat pathogen *Zymoseptoria tritici* triggered cell death in *N. benthamiana* [32]. Of these, 12 required the presence of a signal peptide to elicit a response and, for three of the 12, this response also relied on both *Nb*BAK1 and *Nb*SOBIR1, suggesting that they are recognized by LRR-RLP immune receptors [32]. In another example, four of 85 CE proteins tested from the broad host-range pathogen *Colletotrichum fructicola* triggered cell death in *N. benthamiana* [30]. In this example, the responses were again dependent on the possession of a signal peptide and the presence of *Nb*BAK1. However, in this case, the responses were independent of *Nb*SOBIR1, suggesting that they are instead recognized by LRR-RLK immune receptors [30].

With the abovementioned studies in mind, we set out to determine if a similar high-throughput screening approach could be used in *Nicotiana* species to identify CE proteins of *V. inaequalis* that are recognized by LRR-RLP or LRR-RLK immune receptors. By screening 137 selected CE proteins of *V. inaequalis*, we aimed to identify exogenous sources of resistance that could be deployed with endogenous sources of resistance in apple to confer more sustainable and broad-spectrum resistance to apple scab.

## Results

### Six CE proteins of *V. inaequalis* trigger cell death upon extracellular targeting to the leaf apoplast of *N. benthamiana* and *N. tabacum*

As a starting point to determine whether *Nicotiana* species possess extracellular immune receptors capable of providing resistance to *V. inaequalis* upon transfer to apple, 137 CE proteins from this fungus were screened for their ability to trigger cell death in wild-type (WT) *N. benthamiana* and *N. tabacum* plants using ATTAs (**Table S1**). These CE proteins were selected on the basis that they are small (≤300 amino acid residues in length), are predicted to be secreted, and are encoded by genes that are expressed during infection of apple by *V. inaequalis* (**Table S1**). Further details of the criteria used for CE protein selection can be found in the Materials and Methods section.

As a positive control, cell death was triggered by the INF1 elicitin protein from the potato/tomato pathogen *Phytophthora infestans* [40, 41] in both *N. benthamiana* and *N. tabacum* (**Figure 1**). As a negative control, the pICH86988 empty vector (EV) did not trigger cell death in either of these species (**Figure 1**). In total, six out of the 137 CE proteins triggered cell death in one or both of *N. benthamiana* and *N. tabacum* (**Figure 1**). More specifically, CE28, CE74 and CE77 triggered cell death in *N. benthamiana* but not *N. tabacum*, while CE33, CE89 and CE128 triggered cell death in both *Nicotiana* species (**Figure 1**). Cell death was dependent on extracellular targeting of the CE proteins to the leaf apoplast, since no cell death was observed in the absence of a signal peptide in *N. benthamiana* (**Figure 1**). This suggests that, if the observed cell death responses are the result of recognition by immune receptors in *N. benthamiana* and *N. tabacum*, these immune receptors are extracellular. However, it must be pointed out that, as each of the six CE proteins lacking a signal peptide did not transition through the endoplasmic reticulum (ER)-Golgi secretory pathway, they would not have undergone post-translational modifications, such as disulphide bond formation, which is anticipated to be required for stability, function and/or recognition. Of note, all six CE proteins lacking a signal peptide were detected in samples from *N. benthamiana* leaves following ATTAs using Western blotting, confirming that they were produced (**Figure S1**).

**Figure 1.**
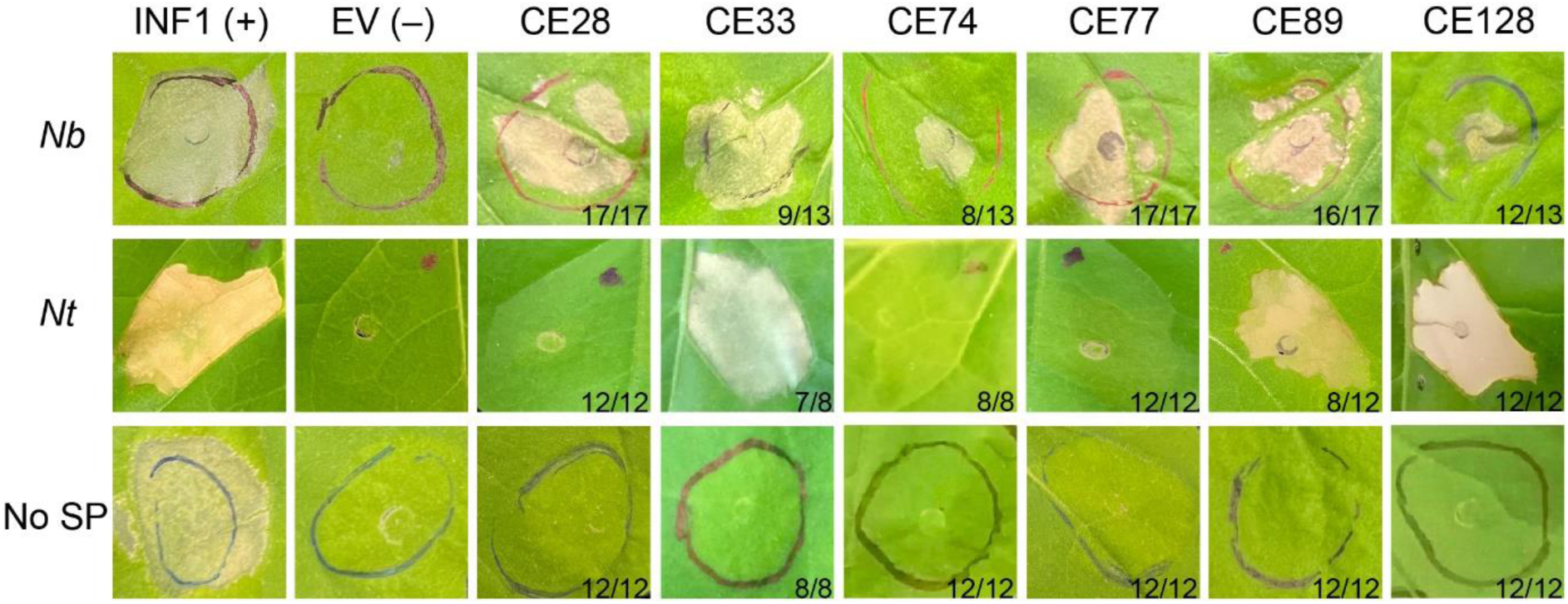
Six candidate effector (CE) proteins of *Venturia inaequalis* trigger cell death upon extracellular targeting to the leaf apoplast of one or both of *Nicotiana benthamiana* and *Nicotiana tabacum*. CE proteins of *V. inaequalis* were expressed in leaves of *N. benthamiana* (*Nb*) and *N. tabacum* (*Nt*) using *Agrobacterium tumefaciens*-mediated transient transformation assays (ATTAs), with each candidate extracellularly targeted to the apoplast using the Pathogenesis-Related 1α (PR1α) signal peptide from *N. tabacum* (*top* and *middle* photos). As a positive (+) control for cell death, the INF1 elicitin from *Phytophthora infestans* was used. As a negative (–) control for cell death, the pICH86988 empty vector (EV) was used. CE proteins were also expressed in *Nb* leaves without the PR1α signal peptide (No SP) using ATTAs (*bottom* photos). Leaves were photographed 6–7 d post-agroinfiltration. Numbers on the bottom right-hand side represent the number of times the response was observed (left) out of the number of times the agroinfiltration was performed (right). The responses/lack of responses observed for control ATTAs were consistently observed across three independent experiments.

A schematic of the six cell death-eliciting CE proteins from *V. inaequalis* is shown in **Figure 2**. Using InterProScan, only three of these proteins are predicted to possess characterized domains. These are CE33, CE74 and CE128, which are predicted to possess an ‘Alternaria alternata a 1’ (AltA1) domain (IPR032382; amino acid residues 37–163), an ‘Egh16-like virulence factor’ (Egh16-like) domain (IPR021476; amino acid residues 22–225), and an ‘expansin_EXLX1’ (Expansin_EXLX1-like) domain (IPR049818; amino acid residues 47–232), respectively (**Figure 2**).

**Figure 2.**
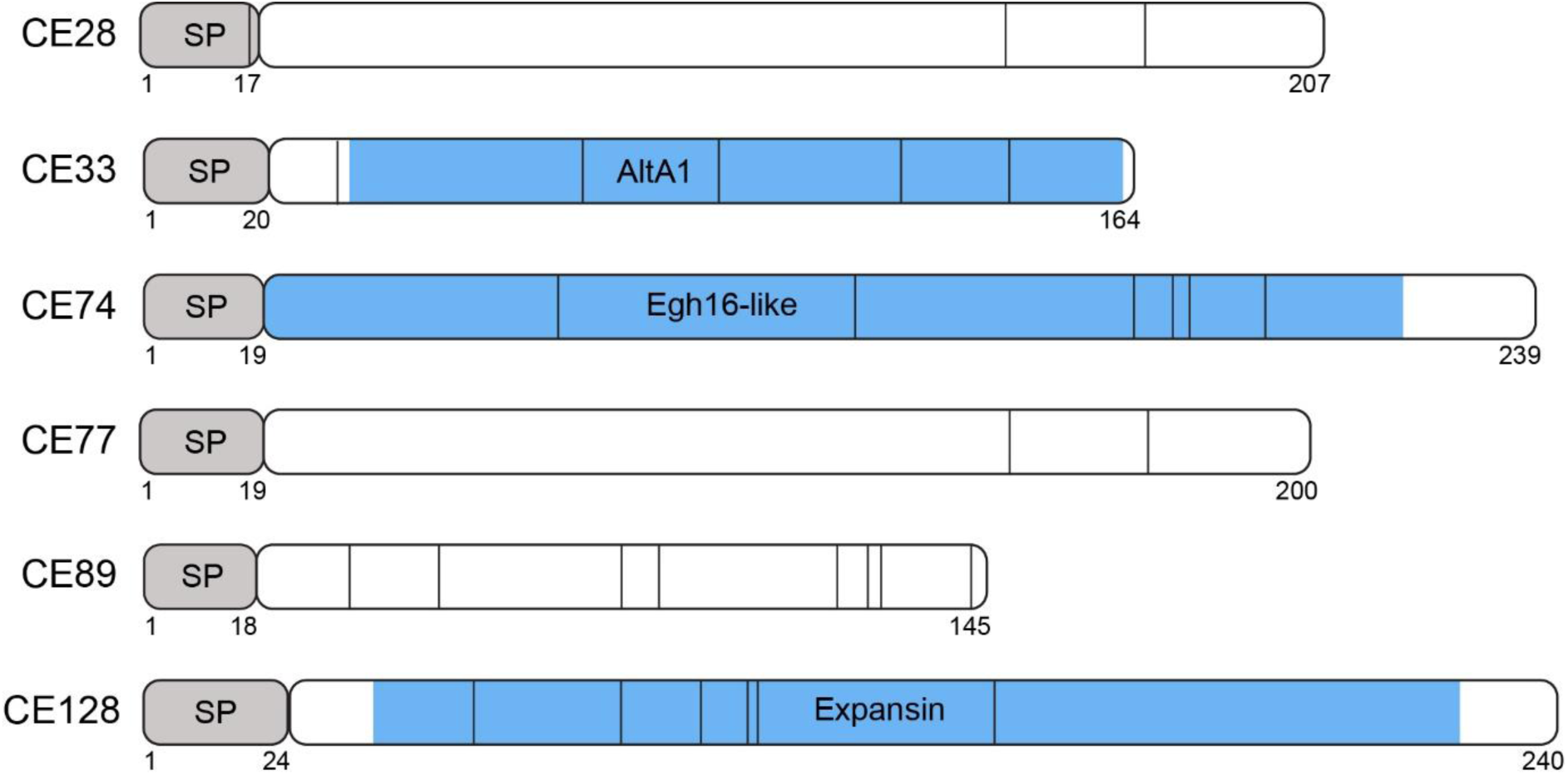
Schematic representation of candidate effector (CE) proteins from *Venturia inaequalis* that trigger cell death in one or both of *Nicotiana benthamiana* and *Nicotiana tabacum*. The signal peptide (SP) of each CE protein for extracellular targeting to the apoplastic environment is shown in grey. Predicted domains are shown in blue and are named accordingly. Cysteine residues are depicted by black vertical lines. Numbers indicate the first and last amino acid residue of each SP and the last residue of each protein.

### Only CE89-mediated cell death is abolished in *ΔNbSOBIR1* and *ΔNbBAK1* plants of *N. benthamiana*

As mentioned above, SOBIR1 and BAK1 are extracellular, membrane-bound co-receptors in plants that interact with RLPs (SOBIR1 and BAK1) and RLKs (BAK1) to ensure transduction of defence response signals following the recognition of effector proteins, PAMPs and DAMPs in the apoplastic environment. To determine whether the cell death responses triggered by the six CE proteins of *V. inaequalis* in *N. benthamiana* are likely to be dependent on extracellular immune receptors, each was expressed in *N. benthamiana* plants lacking *NbSOBIR1* (*ΔNbSOBIR1* [42]) or *NbBAK1* (*ΔNbBAK1* [43]). Here, an Avr9B-like protein from the tomato pathogen *Stemphylium lycopersici*, TW65_01570, which is known to trigger cell death in *N. benthamiana* independent of *Nb*SOBIR1 [44], was used as a positive control, while the Cf-9 RLP resistance protein/Avr9 avirulence effector protein pair from the tomato–*Fulvia fulva* pathosystem [45, 46], which is known to trigger cell death in WT but not *ΔNbSOBIR1* and *ΔNbBAK1* plants of *N. benthamiana,* was used as a positive (WT plants) and negative (*ΔNbSOBIR1* and *ΔNbBAK1* plants) control.

As expected, TW65_01570 triggered cell death in WT, *ΔNbSOBIR1* and *ΔNbBAK1* plants, whereas the Cf-9/Avr9 pair only triggered cell death in WT plants (**Figure 3**). For CE89, the cell death response was abolished in *ΔNbSOBIR1* and *ΔNbBAK1* plants (**Figure 3**), indicating that this CE protein is recognized by an extracellular LRR-RLP immune receptor in *N. benthamiana*. For CE28, CE33, CE74, CE77 and CE128, a cell death response was still observed in *ΔNbSOBIR1* and *ΔNbBAK1* plants (**Figure 3**), demonstrating that these responses are not dependent on *Nb*SOBIR1 or *Nb*BAK1.

**Figure 3.**
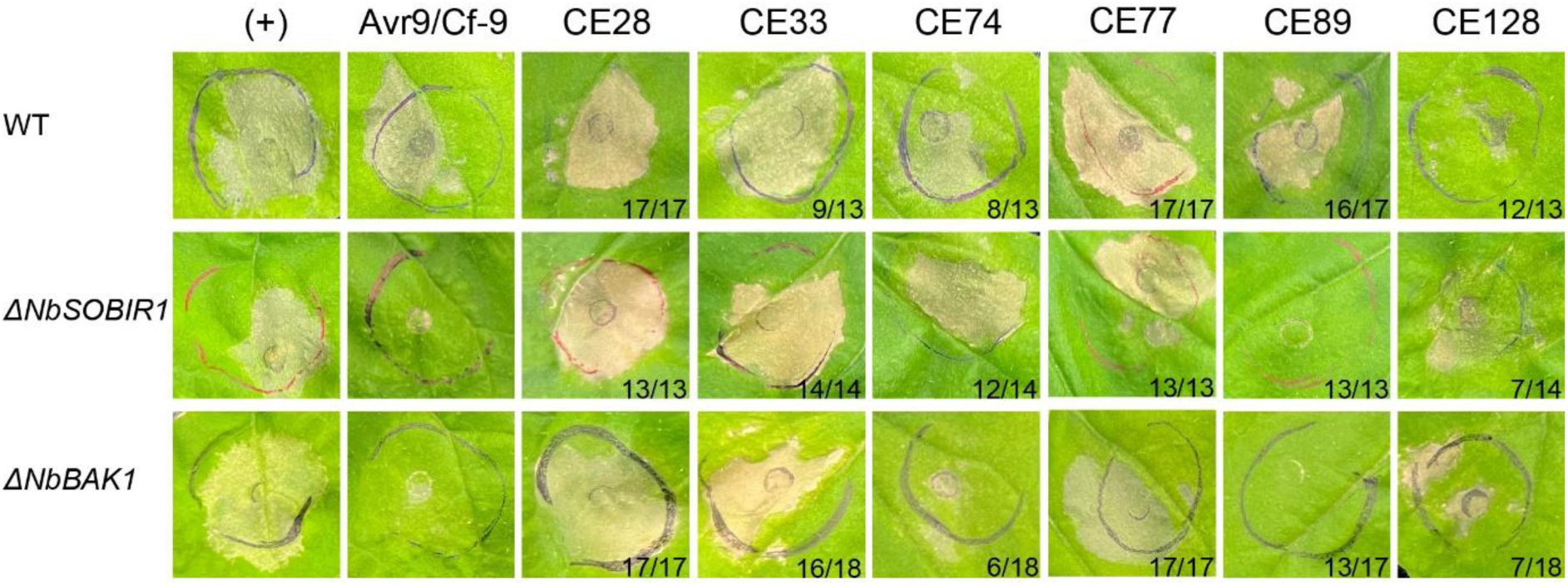
Most cell death responses triggered by candidate effector (CE) proteins of *Venturia inaequalis* in *Nicotiana benthamiana* are independent of *Nb*SOBIR1 and *Nb*BAK1. The CE proteins CE28, CE33, CE74, CE77, CE89 and CE128 of *V. inaequalis* were expressed in leaves of wild-type (WT) *N. benthamiana* or *N. benthamiana* mutants lacking either *SOBIR1* (*ΔNbSOBIR1*) or *BAK1* (*ΔNbBAK1*) using *Agrobacterium tumefaciens*-mediated transient transformation assays (ATTAs), with each candidate extracellularly targeted to the apoplast using the Pathogenesis-Related 1α (PR1α) signal peptide from *Nicotiana tabacum*. As a positive (+) control for cell death across WT, *ΔNbSOBIR1* and *ΔNbBAK1* plants, the Avr9B-like protein from *Stemphylium lycopersici*, TW65_01570, was used. Here, the Cf-9 RLP resistance protein/Avr9 avirulence effector protein pair from the tomato–*Fulvia fulva* pathosystem, also served as a positive control in WT plants. As a negative control for cell death in *ΔNbSOBIR1* and *ΔNbBAK1* plants, the Cf-9/Avr9 RLP resistance protein/avirulence effector protein pair was used. Leaves were photographed 6 d post-agroinfiltration. Numbers on the bottom right-hand side represent the number of times the response was observed (left) out of the number of times the agroinfiltration was performed (right). The responses/lack of responses observed for control ATTAs were consistently observed across three independent experiments.

### Cell death-eliciting CE proteins show limited sequence variation across *V. inaequalis* isolates

To assess the extent of allelic variation in cell death-eliciting CE proteins across *V. inaequalis* isolates pathogenic on *M.* × *domestica*, each was identified in the 77 publicly available *V. inaequalis* genomes in the NCBI database using a tBLASTn search (**Table S2**). The alleles used in the ATTAs were cloned from strains MNH120 or EU-B04 and are designated as the “A” allele for each CE (**Figure S2**). For CE33, CE74, CE89 and CE128, the cloned alleles were the most widely distributed allelic variants found in global *V. inaequalis* populations, whereas for CE28 and CE77, they were the second most abundant alleles. Overall, the six CE proteins are highly conserved across the isolates for which genomes are available. Notably, for CE33, CE77, CE89 and CE128, many of the observed polymorphisms fall within the signal peptide region and are therefore unlikely to affect the mature secreted protein. All signal peptide variants remained predicted as functional by SignalP v4.1 [47]. Taken together, these analyses confirm that the six cell death-eliciting CE proteins show very strong conservation across *V. inaequalis* isolates, consistent with a possibly evolutionary constrained role during infection.

### Cell death-eliciting CE proteins of *V. inaequalis* have sequence and predicted structural similarity to known cell death elicitors from other filamentous plant pathogens

To determine whether the cell death-eliciting CE proteins of *V. inaequalis* have similarity to known cell death elicitors from other filamentous plant pathogens, each was screened against the non-redundant (nr) protein database at the National Center for Biotechnology Information (NCBI) using BLASTp. At the same time, the characterized domain names assigned to CE33 (Alta1/Alt a 1), CE74 (Egh16-like) and CE128 (expansin) were used together with the keywords “cell death” in Google Scholar searches to identify additional potential relationships in the published literature. Then, to confirm these relationships, the tertiary structures of the CE proteins, together with the cell death-eliciting proteins identified from other filamentous plant pathogens in BLASTp and Google Scholar searches, were predicted using AlphaFold3 [48] and compared.

Together, these analyses showed that all six cell death-eliciting CE proteins of *V. inaequalis* have similarity to known cell death elicitors from other filamentous plant pathogens. An example of sequence and predicted structural similarity for each CE protein is shown in **Figures 4 and 5**, respectively. In all cases, alignments between the predicted tertiary structures of the CE proteins and the similar proteins identified by the searches revealed the presence of one or more conserved disulphide bonds (**Figure 5**).

**Figure 4.**
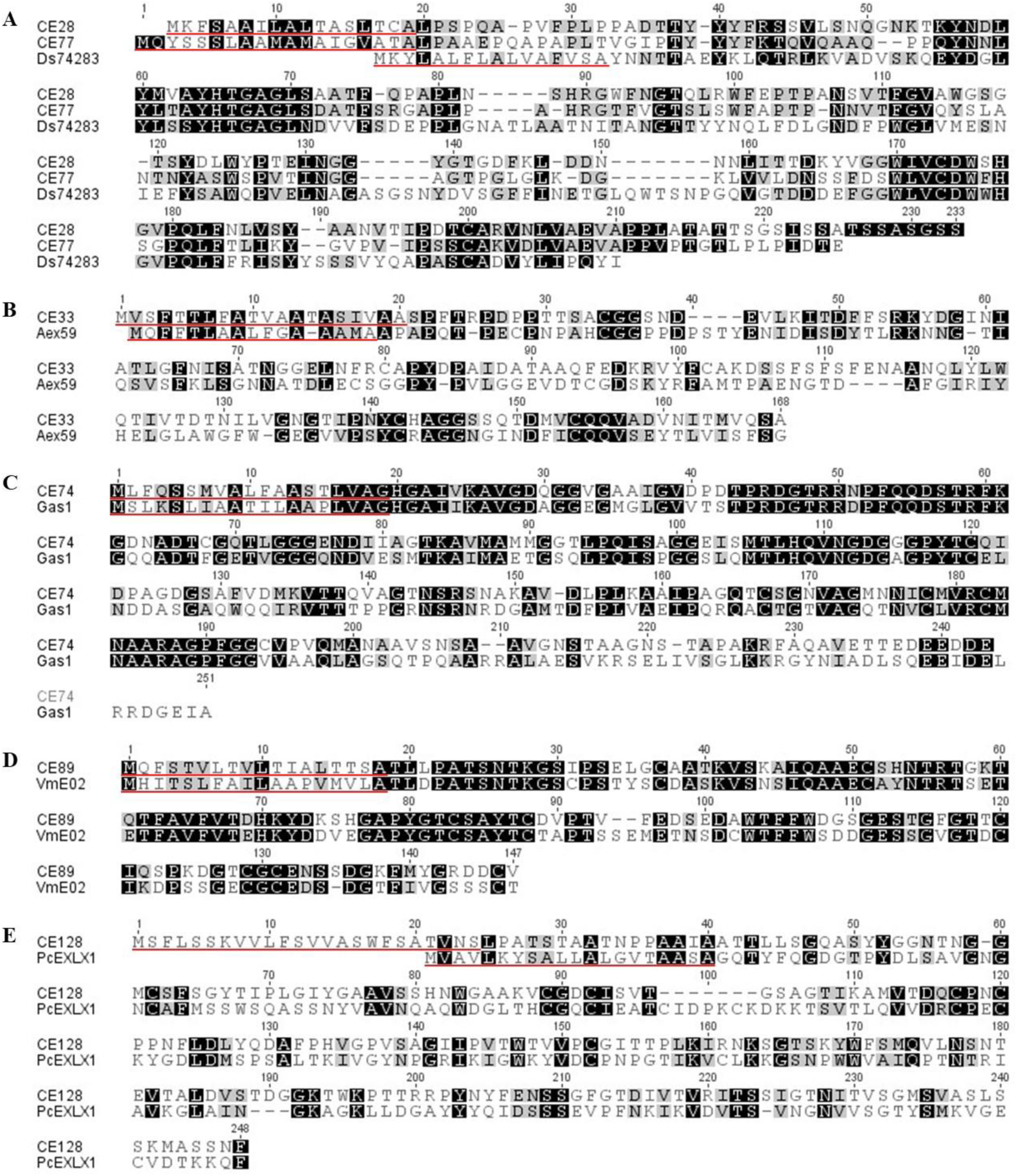
Cell death-eliciting candidate effector (CE) proteins of *Venturia inaequalis* have sequence similarity to known cell death elicitors from other filamentous plant pathogens. **A.** Alignment of CE28 and CE77 to the cell death elicitor *Ds*74283 from *Dothistroma septosporum* (both 29.4% sequence identity) [31, 49]. **B.** Alignment of CE33 to the cell death elicitor Aex59 from *Alternaria solani* (27.5% sequence identity) [50]. **C.** Alignment of CE74 to the cell death elicitor Gas1 from *Magnaporthe oryzae* (52.3% sequence identity) [51, 52]. **D.** Alignment of CE89 to the cell death elicitor *Vm*E02 from *Valsa mali* (57.1% sequence identity) [53]. **E.** Alignment of CE128 to the cell death elicitor *Pc*EXLX1 from *Phytophthora capsici* (23.7% sequence identity) [54]. Identical and similar amino acid residues are highlighted black and grey, respectively. Signal peptide sequences are underlined in red.

**Figure 5.**
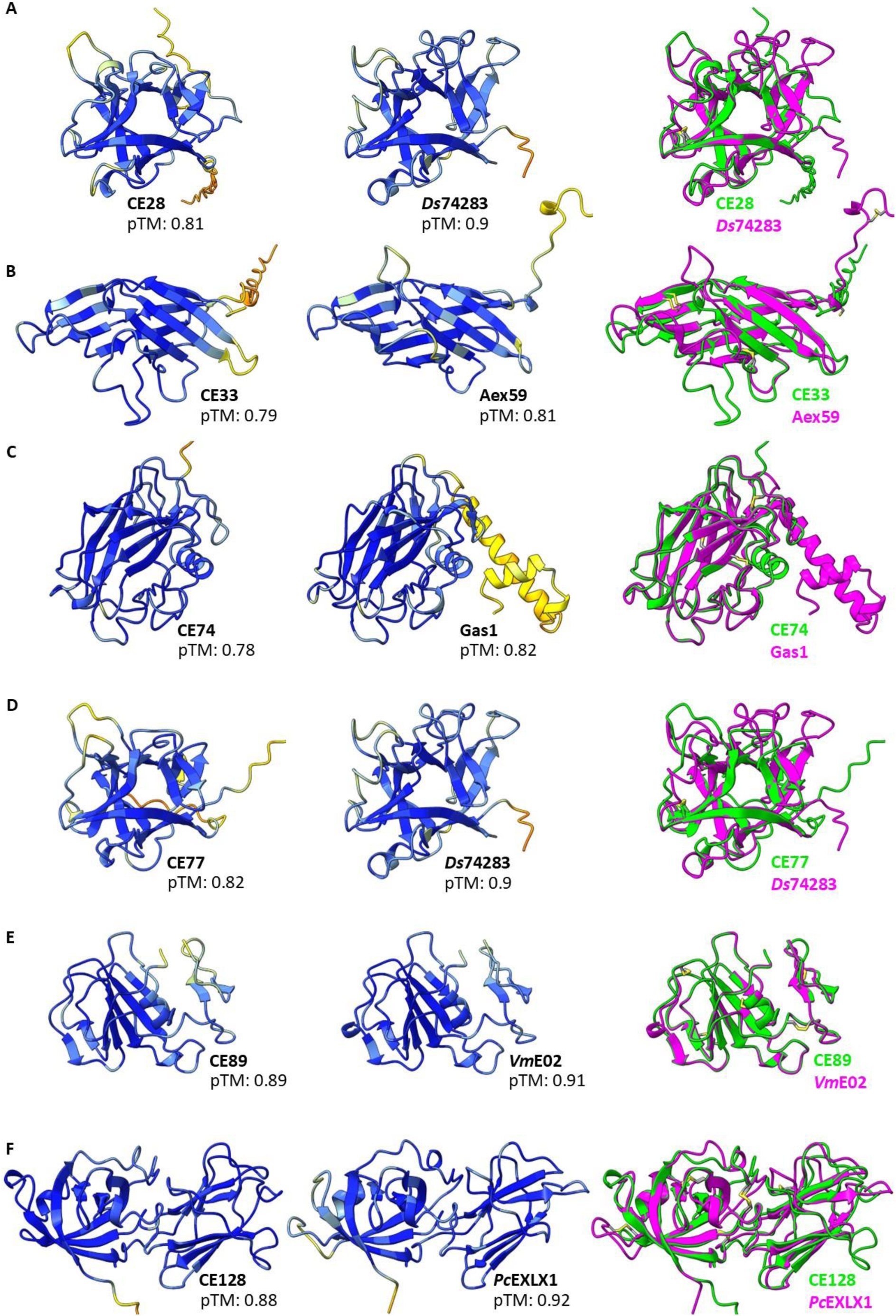
Cell death-eliciting candidate effector (CE) proteins of *Venturia inaequalis* have predicted structural similarity to known cell death elicitors from other filamentous plant pathogens. Predicted tertiary structures of **A.** CE28 (*left*) and the *Dothistroma septosporum* elicitor *Ds*74283 [31, 49] (*middle*), **B.** CE33 (*left*) and the *Alternaria solani* elicitor Aex59 [50] (*middle*), **C.** CE74 (*left*) and the *Magnaporthe oryzae* elicitor Gas1 [51, 52] (*middle*), **D.** CE77 (*left*) and the *D. septosporum* elicitor *Ds*74283 [31, 49] (*middle*), **E.** CE89 (*left*) and the *Valsa mali* elicitor *Vm*E02 [53] (*middle*), and **F.** CE128 (*left*) and the *Phytophthora capsici* elicitor *Pc*EXLX1 [54] (*middle*). Tertiary structures are colour-coded according to AlphaFold predicted local distance difference test (pLDDT) confidence scores (orange = very low, yellow = low, light blue = high, dark blue = very high). pTM: predicted Template Modelling score. In **A–F**, alignments of the CE proteins with the known elicitors are shown on the *right*, with disulphide bonds coloured yellow. RMSD values from structural alignments are as follows for pruned and total atom pairs: CE28 vs *Ds*74283: 0.810 Å (112 pairs) and 3.702 Å (151 pairs); CE33 vs Aex59: 1.013 Å (68 pairs) and 14.316 Å (131 pairs); CE74 vs Gas1: 0.677 Å (165 pairs) and 14.812 Å (189 pairs); CE77 vs *Ds*74283: 0.791 Å (105 pairs) and 3.855 Å (149 pairs); CE89 vs *Vm*E02: 0.388 Å (123 pairs) and 0.611 Å (126 pairs); CE128 vs *Pc*EXLX1: 0.915 Å (135 pairs) and 3.642 Å (195 pairs). Terminal intrinsically disordered regions (IDRs) in the predicted tertiary structures of CE74, CE77, CE128 and Aex59 are excluded for simplicity.

### Cell death-eliciting CE proteins of *V. inaequalis* are encoded by genes that are expressed during infection of apple

To more closely examine the expression profile of the genes encoding the cell death-eliciting proteins from *V. inaequalis*, each was queried against pre-existing RNA-sequencing (RNA-seq) data from Rocafort *et al*. [19]. These data are based on an extensive infection time course of *V. inaequalis* isolate MNH120 on leaves of susceptible apple cultivar ‘Royal Gala’ at 12-and 24-h post-inoculation (hpi), as well as 2, 3, 5 and 7 d post-inoculation (dpi), and in culture on the surface of cellophane membranes overlaying potato dextrose agar at 7 dpi, each with four biological replicates. This analysis revealed that *CE28* is most highly expressed in culture and lowly expressed *in planta* (**Figure 6**). *CE33*, on the other hand, although most highly expressed in culture, is also highly expressed throughout *in planta* growth (**Figure 6**). Unlike *CE28* and *CE33*, both *CE74* and *CE77* are expressed at a negligible level in culture but are moderately and highly expressed *in planta*, respectively, with peak expression observed at 12 hpi–2 dpi (*CE74*) and 3–5 dpi (*CE77*) (**Figure 6**). Finally, *CE89* and *CE128* are moderately expressed both in culture and across all *in planta* time points (**Figure 6**).

**Figure 6.**
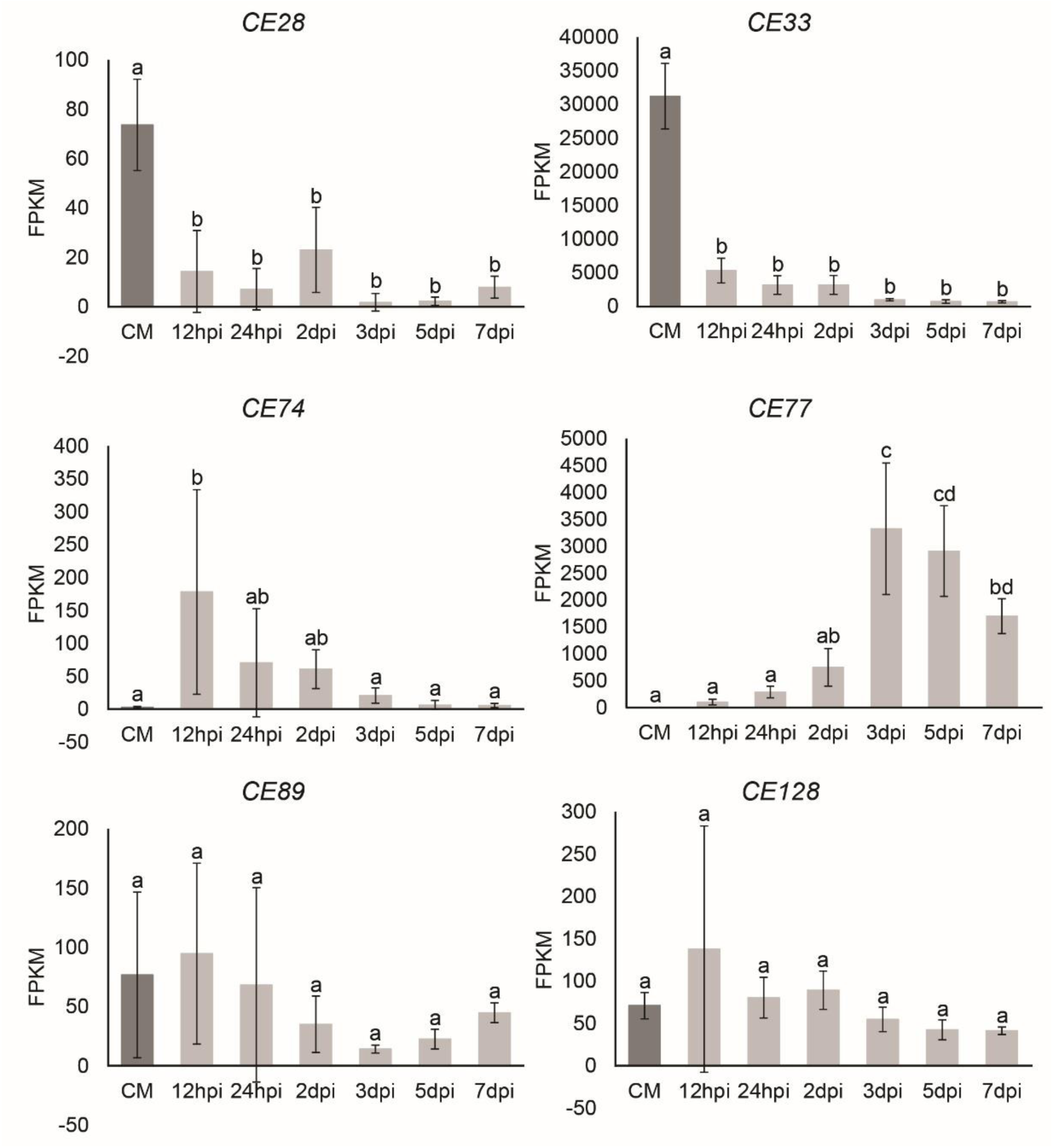
Expression profiles of genes encoding cell death-eliciting candidate effector (CE) proteins of *Venturia inaequalis*. Gene expression levels, shown as RNA-sequencing (RNA-seq) fragments per kilobase of transcript per million mapped reads (FPKM) values, are from an infection time course of *V. inaequalis* isolate MNH120 on leaves of the susceptible apple cultivar ‘Royal Gala’. Sampling was performed at 12 and 24 hours post-inoculation (hpi), as well as 2, 3, 5 and 7 days post-inoculation (dpi; light grey bars) [19]. Expression in culture was measured for the same isolate grown for 7 days on the surface of cellophane membranes (CM) overlaying potato dextrose agar (dark grey bars) [19]. FPKM means and standard error were calculated from four biological replicates. Results were statistically analysed by a one-way ANOVA with *post hoc* Tukey-B test at the 95% confidence level.

### CE33 triggers cell death following infiltration into apple leaves

As shown above, the CE33 protein of *V. inaequalis* is encoded by a gene that, when considered alongside the other five genes that encode cell death-eliciting proteins of this fungus, is by far the most highly expressed during early infection of apple. Whilst CE33 triggers a cell death response that is independent of *Nb*SOBIR1 and *Nb*BAK1 in *N. benthamiana*, it remains possible that other, non-LRR-RLP/LRR-RLK-type extracellular immune receptors in this plant species are responsible for its recognition. Such an immune receptor, which has the potential to prevent ingress of *V. inaequalis* during early infection, is of high value for durable disease resistance against scab disease. Before this potential source of resistance can be investigated further, we set out to determine whether CE33 also triggers cell death in apple. If so, this may suggest that apple already has an immune receptor capable of recognizing CE33.

To investigate whether CE33 triggers cell death in apple, we first heterologously produced allele “A” of this protein in *Pichia pastoris* and purified it for leaf infiltration assays (**Figure S3**). To confirm that the purified CE33 protein was active, it was infiltrated into leaves of *N. tabacum* and compared with elution buffer alone. As expected, the buffer did not induce cell death, while CE33 triggered a strong response, confirming its activity ( **Figure 6**). Similarly, in all apple cultivars tested, including ‘Royal Gala’ selection ‘Galaxy’ ( lacking any characterized *Rvi* genes), as well as accessions TSR33T239/X2249 (carrying *Rvi4*) and OR45T132 (carrying *Rvi5*), the elution buffer alone did not cause cell death (**Figure 7**). In contrast, CE33, tested at a concentration of 25 µM, triggered a cell death response in all apple genotypes tested (**Figure 7**). Notably, in both *N. tabacum* and apple, the response was rapid, with cell death visible within 4 h. These results suggest that, at the tested concentration, apple has the capacity to recognize CE33 or, alternatively, that CE33 acts as a non-specific elicitor or toxin capable of activating cell death independently of immune receptor recognition.

**Figure 7.**
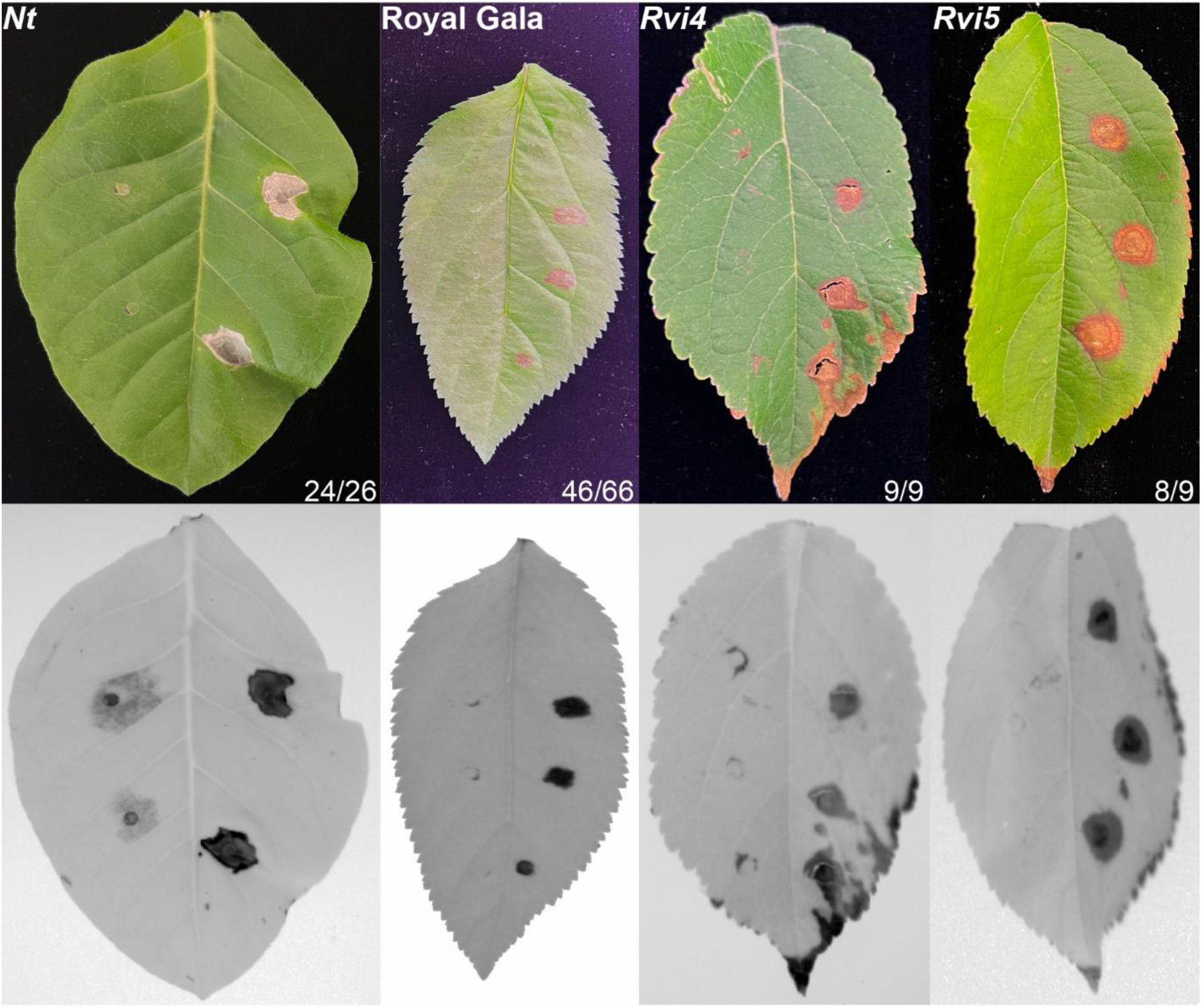
The CE33 protein of *Venturia inaequalis* triggers cell death in *Nicotiana tabacum* (*Nt*) and apple. Purified CE33 protein (*right*) at a concentration of 25 µM was infiltrated into the leaves of *N. tabacum*, apple cultivar ‘Royal Gala’ (no *Rvi* genes), or apple accessions TSR33T239/X2249 (carrying the *Rvi4* gene) and OR45T132 (carrying the *Rvi5* gene) and compared with infiltrated protein buffer alone (negative control; *left*). Photographs with visible light (*top*) and infra-red light (*bottom*) were taken 4 d post-infiltration. Numbers on the bottom right represent the number of times the response was observed (*left*) out of the number of times the infiltration was performed (*right*).

## Discussion

This study aimed to identify CE proteins of *V. inaequalis* that are recognized by extracellular immune receptors of the model plant species *N. benthamiana* and *N. tabacum* using ATTAs. The rationale behind this approach was that the exogenous resistance genes encoding these immune receptors have not co-evolved with *V. inaequalis*, in contrast to the endogenous *Rvi* genes that have interacted for millennia with populations of this fungus on wild *Malus* species in natural environments [11], and are therefore expected to confer more durable resistance against scab disease. Once identified, these exogenous resistance genes could be stacked together with endogenous *Rvi* genes in apple cultivars by genetic modification.

For the ATTAs, a cell death response was used as a visual indicator of potential recognition by an extracellular immune receptor upon targeting of the CE protein to the leaf apoplast. In total, three CE proteins exclusively triggered cell death in *N. benthamiana* (**Table S3**), of which one, CE74, has an ‘Egh16-like virulence factor’ (Egh16-like) domain. Effector proteins with this domain predominantly play important roles during early host colonization, contributing to host penetration and suppression of host immunity [52, 55–57]. Consistent with this role, *CE74* is most highly expressed from 12 hpi to 2 dpi, when *V. inaequalis* is forming appressoria on the apple leaf surface and stromata undeath the host cuticle [19]. To the best of our knowledge, CE74 is the first Egh16-like protein identified to date that triggers cell death in *N. benthamiana*, although Gas1 from *M. oryzae* triggers cell death upon transient expression in rice protoplasts [51]. However, whether Gas1 is a secreted protein remains unclear, despite being predicted to possess a signal peptide [52]. For CE74, the cell death response triggered in *N. benthamiana* occurred independently of *Nb*SOBIR1 and *Nb*BAK1, suggesting that this protein is not recognized by an LRR-RLP or LRR-RLK immune receptor.

Two additional CE proteins of *V. inaequalis* that exclusively triggered cell death in *N. benthamiana* were CE28 and CE77. Of these, CE77 is encoded by a gene that is highly expressed at 3–5 dpi, when infection structures of *V. inaequalis* are rapidly expanding in size and number beneath the apple cuticle [19]. In contrast, CE28 is encoded by a gene that is poorly expressed *in planta*, instead peaking in culture. Both proteins adopt a β-trefoil fold and share sequence and/or structural similarity with *Ds*74283 and *Ds*71487 from the pine pathogen *Dothistroma septosporum*, which trigger cell death in pine and *N. benthamiana* [31, 49, 58]. *Ds*71487 itself is a sequence homolog of *Rc*CDI1 from the barley pathogen *Rhynchosporium commune*, which induces cell death in *N. benthamiana* and other members of the Solanaceae [59]. Additional *Rc*CDI1 homologs from *M. oryzae* and *Z. tritici*, as well as the broad host-range pathogens *Botrytis cinerea* and *Sclerotinia sclerotiorum*, also trigger cell death in *N. benthamiana* [59], as do structurally related members of the Ecp32 family from *D. septosporum*, the tomato pathogen *F. fulva*, and the pine pathogen *Cyclaneusma minus* [31, 49, 60, 61]. Despite this common activity, the biological functions of β-trefoil-type effector proteins remain unknown. Notably, the cell death response triggered by CE28 and CE77 occurred independently of *Nb*SOBIR1 and *Nb*BAK1, indicating that these proteins are likely not recognized by an LRR-RLP or LRR-RLK immune receptor in *N. benthamiana*. A similar observation has been made for *Ds*74283, *Ds*Ecp32-1 and *Ds*Ecp32-3 of *D. septosporum*, which also trigger cell death in *N. benthamiana* independently of these co-receptors [49, 61]. In contrast, the cell death triggered by *Ff*Ecp32-3 of *F. fulva* and *Rc*CDI1 of *R. commune* in *N. benthamiana* is dependent on *Nb*SOBIR1 and *Nb*BAK1 [59, 61]. Together, these findings suggest that some β-trefoil-type effector proteins are perceived by an LRR-RLP immune receptor, while others are not.

A further three CE proteins of *V. inaequalis* triggered cell death in both *N. benthamiana* and *N. tabacum*, including CE89 (**Table S3**). CE89 shows sequence and structural similarity to *Vm*E02 of *Valsa mali*, which itself triggers cell death in *N. benthamiana*, tomato, pepper, *Arabidopsis thaliana*, wheat, and apple [53]. Homologs of *Vm*E02, including those from *B. cinerea* (*Bc*Plp1**/**SCP*^Bc^*), *S. sclerotiorum* (SCP*^Ss^*), *C. minus* (*Cm*2721), *D. septosporum* (*Ds*131885), the wheat pathogen *Fusarium pseudograminearum* (*Fp*00392), and the pine pathogen *Phytophthora pluvialis* (*Pp*10632), also trigger cell death in one or both *Nicotiana* species [49, 53, 62]. Consistent with CE89, the *Vm*E02-induced cell death response in *N. benthamiana* is dependent on *Nb*SOBIR1 and *Nb*BAK1 [53], while for *Cm*2721, *Ds*131885 and *Pp*10632, a requirement for *Nb*SOBIR1 has been shown [49]. In contrast, the cell death response triggered by *Fp*00392 in *N. benthamiana* is not dependent on either co-receptor [62]. As the biological functions of *Vm*E02-type proteins differ among species, with some contributing to asexual reproduction, stress tolerance or virulence [53, 62], the role of CE89 during growth in culture or host infection remains unclear.

In *N. benthamiana*, *Vm*E02 triggers cell death upon recognition by the RLP RE02 [53, 63], while in *A. thaliana*, SCP*^Ss^* of *S. sclerotiorum* activates immune responses upon recognition by the RLP RLP30 [64]. Given that RE02 contributes to resistance against pathogens in *N. benthamiana* [63], and that RLP30 recognizes not only SCP*^Ss^*, but also its homologs from *B. cinerea* (*Bc*Plp1**/**SCP*^Bc^*) and *P. infestans* [64], it seems conceivable that these RLPs would provide protection against *V. inaequalis* upon genetic transfer to apple. In support of this, RLP30 perceives SCP*^Ss^* upon genetic transfer to *N. benthamiana* plants silenced for *RE02*, while responsiveness to SCP*^Ss^*in *A. thaliana* is provided by introduction of *RE02* into plants deleted for *RLP30* [64]. Curiously, as *Vm*E02 triggers cell death in apple, there is the possibility that an extracellular immune receptor exists in this species that can recognize this protein and, by association, CE89. However, given the lack of sequence similarity in apple, this putative immune receptor appears to be unrelated to RE02 or RLP30 [63, 64].

The second CE protein of *V. inaequalis* that triggered cell death in both *N. benthamiana* and *N. tabacum* was CE128, which is predicted to possess an ‘Expansin’ domain. To the best of our knowledge, CE128 represents the first expansin-like protein to be identified from a plant-pathogenic fungus that triggers cell death in *Nicotiana* species. Several expansin-like effector proteins have, however, been identified from plant-parasitic nematodes that trigger cell death in one or more plant species, including *N. benthamiana* [65–67]. Furthermore, an expansin-like effector protein from the broad host-range pathogen *Phytophthora capsici*, *Pc*EXLX1, has been identified that triggers cell death in *N. benthamiana* [54]. For *Pc*EXLX1, this response is dependent on *Nb*SOBIR1 and *Nb*BAK1, with its perception regulated by a G-type lectin-RLK *Nb*ERK1 [54]. Consistent with the recognition of *Pc*EXLX1 by an immune receptor, deletion of *PcEXLX1* results in enhanced virulence of *P. capsici* on *N. benthamiana*, while overexpression results in decreased virulence [54]. *Nb*ERK1 is unlikely to also regulate the cell death response to CE128, given that the response triggered by this protein was independent of both *Nb*SOBIR1 and *Nb*BAK1 and, therefore unlikely to be recognized by an LRR-RLP or LRR-RLK immune receptor. In plants, expansins are characterized as non-enzymatic proteins with cell wall-loosening activity [68]. Given that CE128 is expressed throughout host infection, it may contribute to loosening the walls at epidermal cell junctions, thereby facilitating appressorium-mediated entry and subsequent subcuticular colonization.

The third CE protein from *V. inaequalis* that triggered cell death in both *N. benthamiana* and *N. tabacum* was CE33, which adopts an Alt a 1-like fold [69, 70]. Strikingly, Alt a 1 proteins from over 25 plant-pathogenic fungal species have been shown to trigger cell death in one or more plant species to date and, in many cases, these plant species include *N. benthamiana* and *N. tabacum* [28, 31, 50, 61, 71–78]. One of these proteins is *Vi*Aex59 of *V. inaequalis*, which triggers cell death in *N. benthamiana* [50] and is CE33 in the current study. Across fungi, Alt a 1-like proteins have been linked to diverse biological functions spanning asexual development, vegetative growth, and stress tolerance, as well as processes that affect the progression of host infection [50, 71, 72, 76–87]. The abundant production of CE33 during host infection [88], together with its strong expression both in culture and *in planta*, suggests that it may play similar roles in *V. inaequalis*.

For several Alt a 1-like proteins, including *Vm*Hrp1 of *V. mali*, *Ss*NE2 of *S. sclerotiorum*, *Ds*Ecp20-3 of *D. septosporum*, and *Ff*Ecp20-3 of *F. fulva*, the ability to trigger cell death in *N. benthamiana* has been linked to recognition by the *Nb*SOBIR1- and *Nb*BAK1-dependent RLP *Nb*RLP26 [61, 73, 74]. A similar immune receptor, *Md*RLP26, has also been identified in apple that can interact with *Vm*Hrp1, suggesting that apple may be capable of detecting Alt a 1-like proteins [73]. Not all Alt a 1-like proteins, however, trigger an *Nb*SOBIR1- and *Nb*BAK1-dependent cell death response. Such proteins are, therefore, unlikely to be recognized by an LRR-RLP or LRR-RLK immune receptor. In addition to CE33, this includes Aex59 from the tomato/potato pathogen *A. solani* [50] and Hip1 from *B. cinerea* [72] in *N. benthamiana*. Interestingly, of these, Hip1 has been shown to lack cytolytic activity and, as such, does not appear to promote cell death through perturbation of the cell wall or plasma membrane [72]. However, upon induction of cell death, an accumulation of ROS, ethylene, salicylic acid, and jasmonic acid, as well as an associated upregulation of defence-related genes, is observed [72]. Similar responses, which are all typical following recognition of a protein by an LRR-RLP or LRR-RLK immune receptor [53, 59, 63, 64, 73], have been shown following cell death elicitation by Aex59 of *A. solani* [50], the *Vm*E02-like protein Fp00392 of *F. pseudograminearum* [62], and the β-trefoil-type protein *Ds*74283 of *D. septosporum* [49] in *N. benthamiana*. Together with the fact that the cell death triggered by CE33, Aex59, *Fp*00392 and *Ds*74283 only occurs upon their targeting to the apoplast [31, 50, 64], like that observed for the other effector proteins described above [31, 49, 53, 54, 59–61, 65, 67, 71–74, 76, 77], this suggests that CE28, CE33, CE74, CE77 and CE128 may trigger non-canonical extracellular immunity in *N. benthamiana* upon recognition by extracellular *Nb*SOBIR1-/*Nb*BAK1-independent immune receptors. Care is needed over this interpretation, however, as DAMPs may be released from necrotic lesions that are then recognized by adjacent cells to trigger immune responses, as has been shown for pathogen necrosis- and ethylene-inducing peptide 1-like (NLP) effector proteins [89].

With the possibility that CE33 triggers non-canonical extracellular immunity, we tested whether this protein also triggers cell death in apple. Regardless of the apple genotype tested, cell death was observed, indicating that this response is not specific to *Nicotiana* species. Interestingly, CE33 was previously identified in culture filtrate samples of *V. inaequalis* that triggered cell death in apple leaves [90, 91]. Following separation based on size and charge, the most abundant protein in the resulting cell death-active fraction was CE33 [90, 91]. Together with the current study, this suggests that CE33 is the protein in the culture filtrate samples that triggered cell death in apple leaves and is consistent with gene expression data showing that *CE33* is very highly expressed during growth of *V. inaequalis* in culture. In any case, this may indicate that CE33 also triggers non-canonical extracellular immunity in apple. Why this protein does not trigger cell death during host infection by *V. inaequalis*, however, remains unknown, especially given that *CE33* is one of the most highly expressed genes *in planta*. One possibility is that the amount of CE33 protein produced by *V. inaequalis* does not reach the threshold required for cell death induction (i.e. the concentration of protein tested, 25 µM, is not biologically relevant). Another possibility is that *V. inaequalis* produces an effector protein that suppresses cell death triggered by CE33, since this response would be detrimental to the biotrophic lifestyle of *V. inaequalis*. Related to this possibility, it has been shown that an effector protein from *A. solani*, Aex113, is able to suppress the cell death and associated transcriptional upregulation of defence genes triggered by Aex59 in *N. benthamiana* [50]. As apple may have the capacity to recognize *Vm*E02-type proteins [53], the same phenomena may also prevent CE89 recognition during infection by *V. inaequalis*.

Taken together, our study has identified six CE proteins of *V. inaequalis* that are candidates for recognition by extracellular immune receptors in *N. benthamiana* and *N. tabacum*. These findings provide a foundation for identifying and deploying corresponding immune receptors in apple to enhance durable scab resistance. One of these CE proteins (CE89) triggered *Nb*SOBIR1- and *Nb*BAK1-dependent cell death, consistent with recognition by a canonical extracellular LRR-RLP immune receptor (likely RE02), whereas the remaining five (CE28, CE33, CE74, CE77 and CE128) acted independently of these co-receptors. Based on parallels with their homologs, it is possible that these five CE proteins are instead recognized by non-canonical extracellular immune receptors. To determine whether this is indeed the case, future research should examine whether defence-related genes are upregulated upon cell death induction in *N. benthamiana* and, importantly, whether the five CE proteins have cytolytic or other toxin-related activities.

Crucially, an analysis of allelic variation across a large collection of publicly available *V. inaequalis* genomes revealed that all six cell death-eliciting CE proteins are highly conserved globally, with only limited polymorphism detected. This strong conservation is consistent with these CE proteins fulfilling evolutionarily constrained roles during infection and suggests that mutations enabling the pathogen to evade recognition by corresponding immune receptors could carry a substantial fitness cost. Nevertheless, for any identified extracellular immune receptors to be effectively deployed in apple, it will need to be determined whether the expression levels and temporal dynamics of the corresponding CE genes are sufficient to permit perception. Furthermore, to assess whether the resistance provided by the immune receptors is likely to be durable, an assessment of whether the identified CE proteins are crucial to *V. inaequalis* virulence will need to be performed using, for example, CRISPR-Cas9 gene editing, which has recently been established in this fungus [92].

Even if recognition is suppressed by other effector proteins of *V. inaequalis* during apple infection, it may be possible to introduce immune receptors with novel recognition specificities to circumvent this suppression. Likewise, if *Md*RLP26 is not the endogenous immune receptor that recognizes CE33 in apple, it could be engineered to do so. Screening additional CE proteins from *V. inaequalis*, such as homologs of glycoside hydrolase enzymes known to be recognized by extracellular immune receptors in *Nicotiana* and other species [93], may also reveal further sources of resistance for transfer to apple. However, any exogenous or engineered endogenous immune receptor must be carefully evaluated to ensure that it does not inadvertently promote disease by necrotrophic or hemibiotrophic pathogens, since several abovementioned CE protein homologs have been suggested to accelerate cell death during necrotrophy or the biotrophic–necrotrophic switch in hemibiotrophy [49, 50, 59, 61, 71, 72].

## Experimental Procedures

### Plant material

#### *N. benthamiana* and *N. tabacum*

WT *N. benthamiana* and *N. tabacum* cultivar Wisconsin 38, as well as *ΔNbSOBIR1* and *ΔNbBAK1 N. benthamiana* [42, 43], were used in this study. Seeds were surface-sterilized with 70% (v/v) ethanol for 5 min, rinsed with sterile water, and germinated in 9×9×8 cm (0.4 L) TEKU pots (Pöppelmann, Lohne, Germany) containing Daltons Premium Seed Mix (Fruitfed, New Zealand) at 21°C in a controlled growth room with 70% humidity and a 12 h light/12 h dark photoperiod. At 2 weeks post-sowing, individual seedlings were transferred to TEKU pots containing Daltons Outdoor Container Mix (Fruitfed, Auckland, New Zealand) and grown at 21°C with the same photoperiod and light intensity conditions.

#### *M.* × *domestica* (apple)

Seedlings derived from open-pollinated seeds of apple cultivar ‘Royal Gala’, selection ‘Galaxy’ (lacking any characterized *Rvi* genes), as well as grafted trees of apple accessions TSR33T239/X2249 (carrying *Rvi4*) and OR45T132 (carrying *Rvi5*), were used in this study. Seeds were surface-sterilized with 75% (v/v) ethanol for 5 min, rinsed five times with sterile deionized water, and then imbibed in sterile water overnight at room temperature. To initiate rapid seed germination, the seed coat (testa) was subsequently removed with forceps and each seed placed into a 50 mL Falcon tube containing 20 mL of 0.5× Murashige and Skoog (MS)-medium (Duchefa Biochemie, Haarlem, The Netherlands) with 1% (w/v) phytagel (Plant Media, Dublin, Ireland). Here, the white embryo of the seed was positioned downwards, with only half of the seed submerged in the MS-phytogel medium. Seeds were then germinated at 21°C with a 16 h light/8 h dark photoperiod for approximately 7 d with the Falcon tube lids on, and then a further 2–3 d with the Falcon tube lids off. Next, germinated seeds with a radicle of approximately 8–10 cm were transferred to 9×9×8 cm (0.4 L) TEKU pots containing Daltons Outdoor Container Mix and allowed to recover in the dark at 21°C for 24 h. Finally, seedlings were maintained at 21°C with a 16 h light/8 h dark photoperiod until required. Grafted apple trees were planted into 20 cm (4.7 L) pots (IP Plastics, Auckland, New Zealand) containing Daltons Outdoor Container Mix and maintained under natural environmental conditions.

### CE cell death screening assay in *N. benthamiana* and *N. tabacum*

#### CE protein selection

CE proteins were selected from *V. inaequalis* isolate EU-B04, which was collected in Belgium and has overcome *Rvi1*- and *Rvi10*-mediated resistance [94, 95], or isolate MNH120 (ICMP 13258), which was collected in New Zealand and has overcome *Rvi1*-mediated resistance [16, 96]. For selection, CE proteins had to (1) be ≤300 amino acid residues in length, (2) possess a signal peptide for extracellular targeting to the subcuticular environment/apoplast, (3) lack a transmembrane domain for integration into the fungal plasma membrane, and (4) lack a glycosylphosphatidylinositol (GPI) anchor lipid modification site for attachment to the outer surface of the fungal plasma membrane or cell wall. Here, signal peptides were predicted using SignalP v4.1 [47], while the absence of transmembrane domains and GPI anchor lipid modification sites was predicted using DeepTMHMM v1.0 [97] and NetGPI v1.1 [98], respectively. In addition to these selection criteria, CE proteins had to be encoded by genes that were expressed during host colonization. More specifically, CE genes had to have an RNA-seq fragments per kilobase of transcript per million mapped reads (FPKM) value of ≥50 during the interaction between isolate EU-B04 and apple cultivar ‘Royal Gala’ at 2 d post-inoculation (dpi), as based on preexisting RNA-seq data from Sannier *et al*. [11], or during the interaction between isolate MNH120 and apple cultivar ‘Royal Gala’ at 12 hours post -inoculation (hpi), 24 hpi, 2 dpi, 3 dpi, 5 dpi or 7 dpi, as based on RNA-seq data from Rocafort *et al*. [19].

#### Vector construction

CE genes from *V. inaequalis* were synthesized in the pTwist Amp vector by Twist Bioscience (San Francisco, CA, USA) and were flanked with *Bsa*I recognition sites for subsequent cloning into the ATTA expression vector, pICH86988 [99], using Golden Gate technology [100]. In each case, the nucleotide sequence encoding the mature CE protein was fused at its 5’ end to nucleotide sequence encoding the *N. tabacum* Pathogenesis-Related 1α (PR1α) signal peptide for extracellular targeting to the leaf apoplast of *N. benthamiana* and *N. tabacum* [101] and a 3xFLAG epitope tag (3xDYKDDDDK) for detection by western blotting [102]. At the same time, a subset of the CE proteins without the PR1α signal peptide nucleotide sequence were also synthesized in pICH86988 to prevent extracellular targeting to the leaf apoplast. pTWIST vectors containing the CEs were propagated in *Escherichia coli* JM110, which is dam/dcm methylation-deficient, and therefore does not methylate *Bsa*I recognition sites, while ATTA expression vectors were propagated in *E. coli* DH5α (Thermo Fisher Scientific, Waltham, MA, USA [103]). Plasmid vectors were subsequently extracted using an E.Z.N.A. Plasmid DNA Mini Kit I (Omega Bio-tek, Norcross, GA, USA), and their insert sequences confirmed with Sanger sequencing technology (Massey Genome Service, Palmerston North, New Zealand) using the pICH86988-F (5′–AGGACACGCTCGAGTATAAG–3′)/pICH86988-R (5′–CATGCGATCATAGGCTTCTC–3′) primer pair. Finally, ATTA expression vectors were transformed into *A. tumefaciens* GV3101 (GoldBio, St Louis, MO, USA [104]) by electroporation as described by Guo *et al*. [105].

#### ATTAs in *N. benthamiana* and *N. tabacum*

Single colonies of *A. tumefaciens* harbouring an ATTA expression vector were cultured overnight at 28°C and 180 rpm (C10 platform shaker, New Brunswick Scientific, Enfield, CT, USA) in lysogeny broth (LB) supplemented with 50 μg/mL kanamycin, 10 μg/mL rifampicin, and 30 μg/mL gentamycin. Following culturing, bacterial cells were pelleted by centrifugation at 2,500 × g for 5 min and resuspended in 1 mL of infiltration buffer (10 mM MgCl_2_·6H_2_O, 10 mM MES-KOH (Sigma-Aldrich, St. Louis, MO, USA), and 100 μM acetosyringone (Sigma-Aldrich)) to an OD_600_ of 0.5. The resulting suspension was infiltrated into the abaxial surface of *N. benthamiana* and *N. tabacum* leaves using a 1 mL needleless syringe. For each CE or control protein, the presence or absence of a cell death response was assessed at infiltration sites on two leaves from two plants (i.e. four infiltration sites), across a minimum of three independent experiments, at 6–7 d post-infiltration. Leaves were photographed using a Nikon D7000 camera. Infra-red (Cy7) images of each leaf were captured with the ChemiDoc MP Imaging System (Bio-Rad, Hercules, CA, USA).

#### Verification of protein expression and expected molecular weights

A Western blot analysis was performed on total protein extracts of *N. benthamiana* leaves following ATTAs using the protocol of Guo *et al*. [105]. For this purpose, proteins were first resolved by Sodium Dodecyl Sulphate Polyacrylamide Gel Electrophoresis (SDS-PAGE) using 12% resolving and 5% stacking gels and transferred to 0.2 μm polyvinylidene fluoride (PVDF) membranes (Bio-Rad). Detection of 3xFLAG-tagged CE proteins was carried out using SuperSignal West Dura Extended Duration substrate (Thermo Fisher Scientific) in conjunction with a mouse-derived anti-FLAG M2 monoclonal antibody (Sigma-Aldrich) and a chicken-derived anti-mouse secondary antibody (Santa Cruz Biotechnology, Dallas, TX, USA). Membranes were imaged using the Azure c600 Bioanalytical Imaging System (Azure Biosystems, Dublin, CA, USA).

### CE cell death screening assay in apple

#### Protein production and purification

Production and purification of 6×His-tagged CE33 protein was performed according to Tarallo *et al*. [106]. Briefly, the nucleotide sequence encoding the mature CE33 protein (i.e. without its native signal peptide) was amplified using a Polymerase Chain Reaction (PCR) experiment from the CE33 ATTA expression vector with the CE33-F (5′–CCCGGGCCCGACTACAAGGACGACGATGACAAGAGCCCATTCACCCGCC–3′)/CE33-R (5′–GGGAATTCCTAGGCGCTCTGGACCATG–3′) primer pair and cloned behind nucleotide sequence encoding an N-terminal 6×His tag in the pPic9-His6 vector [107] for expression in *P. pastoris* GS115 (Thermo Fisher Scientific). Here, the CE33-F primer incorporated nucleotide sequence encoding a 5′ FLAG-tag for detection by Western blotting. After propagation in *E. coli* DH5α, and sequence confirmation using Sanger sequencing technology, the resulting expression vector was linearized with *Sma*I (incorporated into the vector using the CE33-F primer) and transformed into *P. pastoris*. Protein expression was induced in Buffered Methanol-Complex Medium (BMMY) using 0.5% methanol over 48 h.

Following centrifugation and filter-sterilization, the culture supernatant was collected and adjusted to pH 7.4. Secreted 6×His-tagged CE33 protein was purified from the filtrate using immobilized metal ion affinity chromatography (IMAC) with a Ni Sepharose™ 6 Fast Flow resin (GE Healthcare, Chicago, IL, USA), following the manufacturer’s guidelines. Eluted fractions were pooled, and buffer exchange was performed to remove imidazole from the purified protein solution. Following affinity purification, the elution buffer was replaced with the same buffer lacking imidazole (20 mM sodium phosphate, 0.5 M NaCl; pH 7.4) using Amicon Ultra-15 centrifugal filter units (Merck Millipore) with an appropriate molecular weight cut-off. The sample was centrifuged according to the manufacturer’s instructions, and the retentate was washed multiple times with buffer to ensure complete removal of residual imidazole.

Protein purification was assessed by SDS-PAGE using 4–20% Mini-PROTEAN TGX Stain-Free™ Protein Gels (Bio-Rad), loaded with up to 20 µL of purified protein per well. The purified protein samples were resolved alongside the Precision Plus Protein™ All Blue Prestained Protein Standard (Bio-Rad) for molecular weight estimation. Electrophoresis was carried out under denaturing conditions in 1× Tris-glycine-SDS running buffer (2.5 mM Tris, 1.92 mM glycine, 0.01% SDS; pH 8.3) at a constant current of 30 mA for 40 min. Protein concentration was estimated using a NanoDrop™ One/OneC Microvolume UV-Vis Spectrophotometer (Thermo Fisher Scientific). Absorbance at 280 nm was used to calculate protein concentration, based on the theoretical extinction coefficient (13,200 M^-1^ cm^-1^) and molecular weight (15,574 Da) of the protein.

#### Protein infiltration into apple

Purified 6×His-tagged CE33 protein at a concentration of 25 µM was infiltrated into the abaxial surface of apple and *N. tabacum* leaves using a 1 mL needleless syringe. At the same time, on the opposite side of the major leaf vein, the protein buffer alone (20 mM sodium phosphate, 0.5 M NaCl; pH 7.4) was infiltrated as a negative cell death control. A total of 66 infiltration zones were analysed for apple cultivar ‘Royal Gala’, with three infiltration zones per leaf. Infiltrations were carried out across leaves sampled from four different apple plants. For the Rvi4 and Rvi5 apple accessions, a total of nine infiltration zones were analysed, with three infiltration zones per leaf across three leaves. All infiltrated leaves were collected from the same plant. A total of 26 infiltration zones were analysed for *N. tabacum*. Each infiltration zone corresponded to one independent infiltration, with two infiltration zones per leaf, resulting in 12 leaves infiltrated in total. These leaves were sampled from six different plants, with two leaves infiltrated per plant. All leaves were photographed at 4 d post-infiltration using a Nikon D7000 camera. Infra-red (Cy7) images of each leaf were captured with the ChemiDoc MP Imaging System (Bio-Rad).

### CE protein bioinformatic analyses

CE BLASTp searches were performed against the nr protein sequence database at NCBI. Tertiary structures were predicted using AlphaFold3 [48] and then aligned and visualized using ChimeraX v1.9 [108]. Here, predicted tertiary structures were coloured according to AlphaFold predicted local distance difference test (pLDDT) score using the command: `[color bfactor palette alphafold]`. Protein sequence alignments were generated using Geneious v9.0.5 [109]. Protein domains were predicted in mature protein sequences (i.e. lacking a signal peptide) using InterProScan [110]. For the allelic variation analysis, sequences of the six cell death-eliciting CEs were blasted against the available *V. inaequalis* genomes at the NCBI. Incomplete sequences due to sequencing or assembly errors were removed from the analysis. Protein sequences alignments were generated and edited using Geneious v9.0.5 [109].

### CE gene expression analysis

The expression of genes from isolate EU-B04 of *V. inaequalis* during infection of apple cultivar ‘Royal Gala’ was assessed using pre-existing RNA-seq data from Sannier *et al*. [11]. The expression of genes from isolate MNH120 of *V. inaequalis* during infection of apple cultivar ‘Royal Gala’, as well as during growth in culture on cellophane membranes overlaying PDA, was assessed using pre-existing RNA-seq data from Rocafort *et al*. [19], in association with the gene models from Deng *et al*. [16] and Rocafort *et al*. [19]. These expression data were previously validated by Rocafort *et al*. [111] using real-time quantitative PCR (RT-qPCR).

## Acknowledgements

We thank Kee Hoon Sohn (Seoul National University; formerly Massey University) for providing seeds of WT *N. benthamiana* and *N. tabacum* cultivar Wisconsin 38, Matthieu Joosten (Wageningen University and Research) for providing seeds of *ΔNbBAK1* and *ΔNbSOBIR1 N. benthamiana*, as well as strains of *A. tumefaciens* carrying the Avr9 and Cf-9C ATTA expression constructs, Vincent Bus (The New Zealand Institute for Bioeconomy Science Limited) for providing grafted *Rvi4* and *Rvi5* apple plants, and Jasna Rakonjac (Massey University) for providing *E. coli* strain JM110. Silvia de la Rosa was supported by a Massey University Doctoral Scholarship.

## Author Contributions

CHM, SdlR, JKB, KMP, REB, and BLC designed the research; SdlR, MT, JT, MS, Y-HT, and AMM performed the research; SdlR, MT, and JT carried out the data analysis, collection or interpretation; SdlR, MT, BLC and CHM wrote the manuscript. All authors reviewed the manuscript and approved it for publication.

## Conflicts of Interest

The authors declare no conflicts of interest.

## Data Availability Statement

The data that support the findings of this study are available in the **Supporting Information** of this article.

## Supporting Information

**Figure S1.**
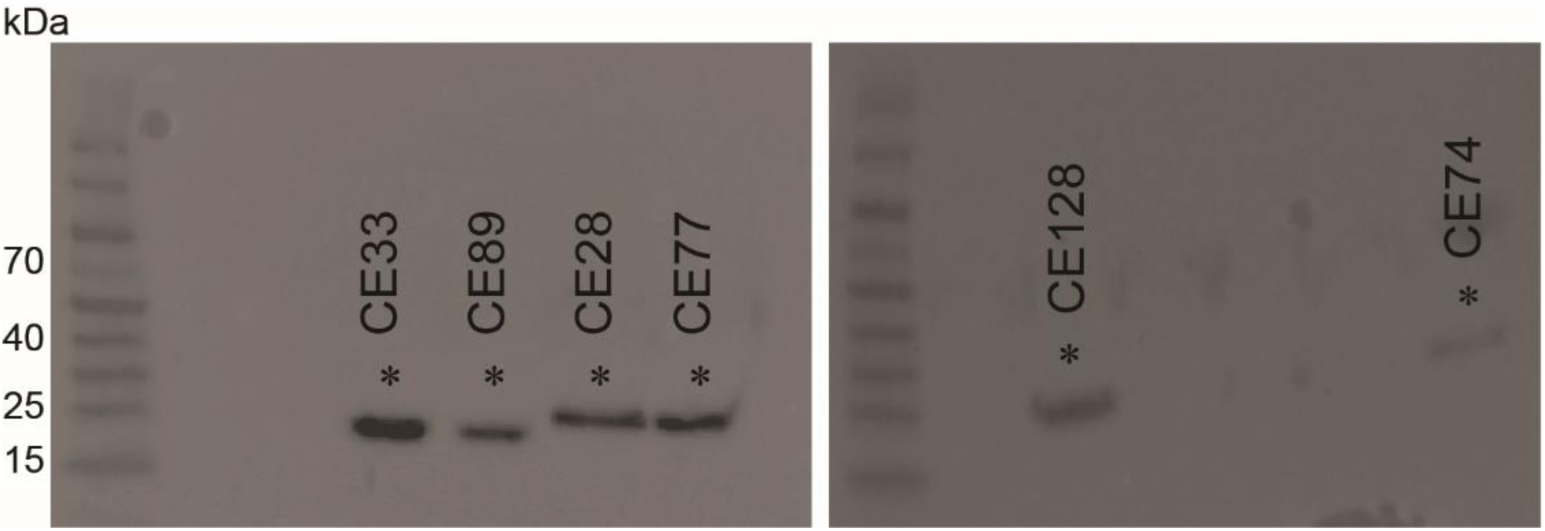
Western blot detection of candidate effector (CE) proteins from *Venturia inaequalis* expressed in *Nicotiana benthamiana*. CE proteins without the Pathogenesis-Related 1α (PR1α) signal peptide from *Nicotiana tabacum* were expressed in *N. benthamiana* leaves using *Agrobacterium tumefaciens*-mediated transient transformation assays (ATTAs). Total protein was extracted from leaves at 2 d post-agroinfiltration and an anti-FLAG antibody used for the immunodetection of CE proteins. The PageRuler Prestained Protein Ladder (Thermo Fisher Scientific) is shown on the left of each membrane. Asterisks (*) have been placed above CE protein bands. The expected sizes of the CE proteins are 20.4 kDa (CE28), 15.5 kDa (CE33), 21.8 kDa (CE74), 19.2 kDa (CE77), 13.5 kDa (CE89), and 22.4 kDa (CE128).

**Figure S2.**
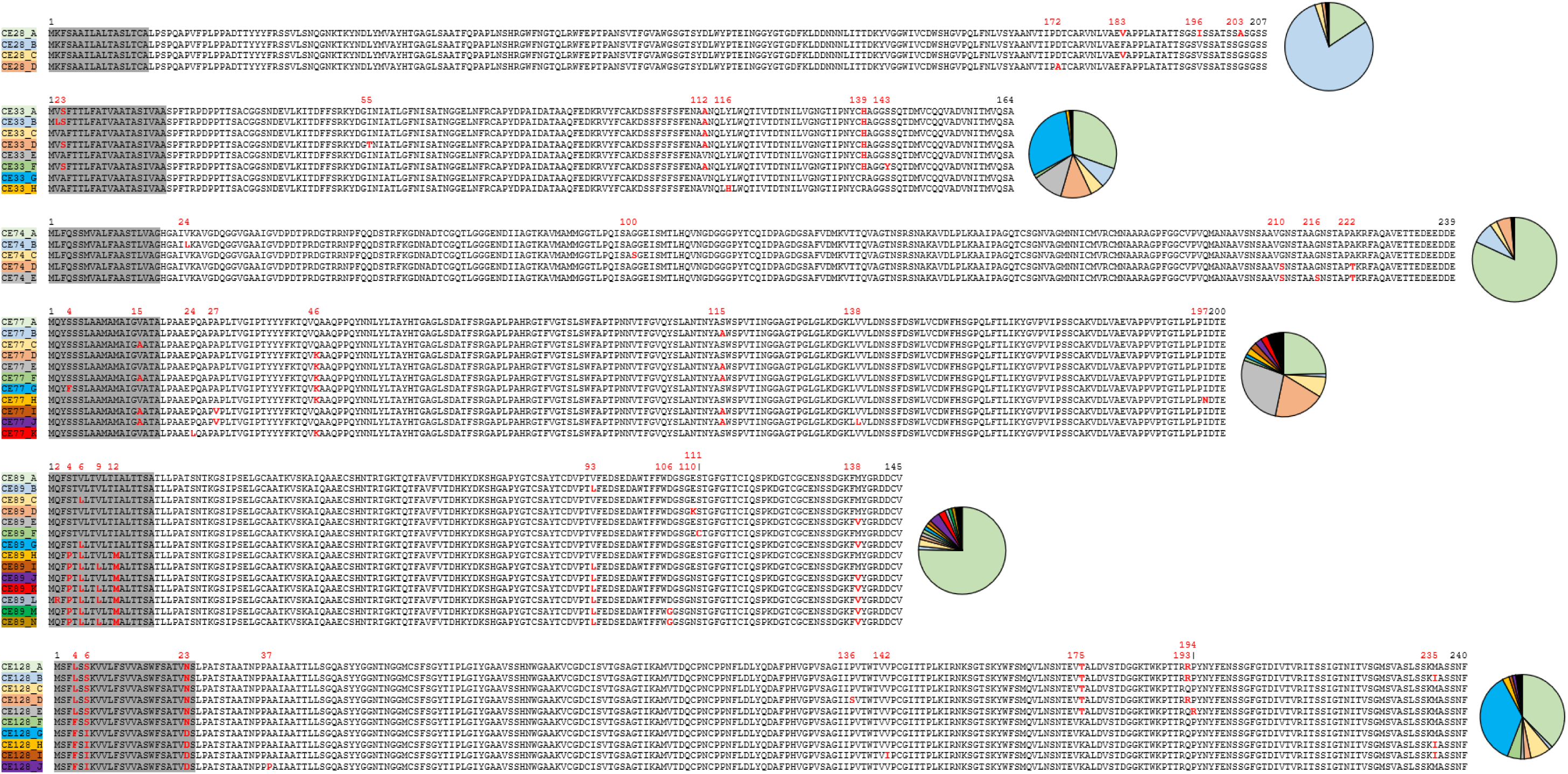
Protein alignments of cell death-eliciting candidate effector (CE) proteins from *Venturia inaequalis*. Signal peptides are highlighted in grey; polymorphisms are colored in red. Pie charts represent the proportion of each allele across 77 *V. inaequalis* isolates pathogenic on *Malus × domestica*.

**Figure S3.**
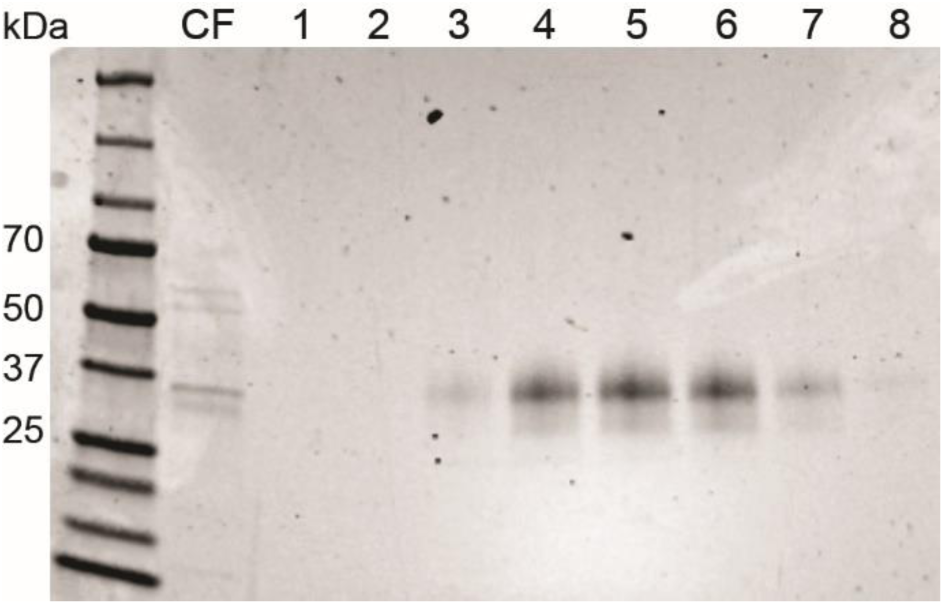
Sodium Dodecyl Sulphate Polyacrylamide Gel Electrophoresis (SDS-PAGE) detection of the purified candidate effector protein, CE33, from *Venturia inaequalis*. Lanes from left to right contain collected fractions during immobilized metal affinity chromatography. *Lane CF:* culture filtrate prior to purification. *Lanes 1–2:* flow-through fractions collected before the elution peak. *Lanes 3–8:* eluted fractions corresponding to the main protein peak, molecular weight markers (kDa) are shown on the left.

**Table S3.** Overview of cell death responses triggered by candidate effector (CE) proteins of *Venturia inaequalis* in leaves of *Nicotiana benthamiana* (*Nb*), *Nicotiana tabacum* (*Nt*), and apple.

| CE protein | Domain/fold | Cell death triggered in Nb? | Cell death triggered in Nt? | Cell death triggered in apple? | Cell death dependent on a signal peptide (SP) in Nb? | Cell death dependent on NbSOBIR1 in Nb? | Cell death dependent on NbBAK1 in Nb? |
| --- | --- | --- | --- | --- | --- | --- | --- |
| CE28 | β-trefoil | Yes | No | Not tested | Yes | No | No |
| CE33 | AltA1-like | Yes | Yes | Yes | Yes | No | No |
| CE74 | Egh16-like | Yes | No | Not tested | Yes | No | No |
| CE77 | β-trefoil | Yes | No | Not tested | Yes | No | No |
| CE89 | VmE02-like | Yes | Yes | Not tested | Yes | Yes | Yes |
| CE128 | Expansin_EXLX1-like | Yes | Yes | Not tested | Yes | No | No |

## Notes

### Competing Interest Statement

The authors have declared no competing interest.

## References

1. Bowen JK, Mesarich CH, Bus VGM, Beresford RM, Plummer KM, Templeton MD. *Venturia inaequalis*: the causal agent of apple scab. Mol Plant Pathol. 2011;12(2):105–22. doi: 10.1111/j.1364-3703.2010.00656.x.

2. Antal G, Szabó S, Szarvas P, Holb IJ. Yield and cost–benefit analyses for apple scab sanitation practices in integrated and organic apple management systems. Plants People Planet. 2024;6(2):470–89. doi: 10.1002/ppp3.10460.

3. MacHardy WE. Apple scab: biology, epidemiology, and management. 1996. pp. xvi+–545.

4. Nasonov AI, Yakuba GV. Apple scab: Resistance to chemical fungicides. Biol Bull. 2024;14(1):S17–S30. doi: 10.1134/S2079086424600802.

5. Bus VGM, Rikkerink EHA, Caffier V, Durel C-E, Plummer KM. Revision of the nomenclature of the differential host-pathogen interactions of *Venturia inaequalis* and *Malus*. Annu Rev Phytopathol. 2011;49:391–413. doi: 10.1146/annurev-phyto-072910-095339.

6. Khajuria YP, Kaul S, Wani AA, Dhar MK. Genetics of resistance in apple against *Venturia inaequalis* (Wint.) Cke. Tree Genet Genomes. 2018;14(2):16. doi: 10.1007/s11295-018-1226-4.

7. Peil A, Howard NP, Bühlmann-Schütz S, Hiller I, Schouten H, Flachowsky H, et al. *Rvi4* and *Rvi15* are the same apple scab resistance genes. Mol Breed. 2023;43(10):74. doi: 10.1007/s11032-023-01421-0.

8. Schouten HJ, Brinkhuis J, van der Burgh A, Schaart JG, Groenwold R, Broggini GAL, et al. Cloning and functional characterization of the *Rvi15* (*Vr2*) gene for apple scab resistance. Tree Genet Genomes. 2014;10(2):251–60. doi: 10.1007/s11295-013-0678-9.

9. Belfanti E, Silfverberg-Dilworth E, Tartarini S, Patocchi A, Barbieri M, Zhu J, et al. The *HcrVf2* gene from a wild apple confers scab resistance to a transgenic cultivated variety. Proc Natl Acad Sci USA. 2004;101(3):886–90. doi: 10.1073/pnas.0304808101.

10. Yousaf A, Baldi P, Piazza S, Gualandri V, Komjanc M, Dalla Costa L, et al. The Hansen’s baccata #2 gene *Rvi12_Cd5* confers scab resistance to the susceptible apple cultivar “Gala Galaxy”. Plant J. 2025;121(2):e17214. doi: 10.1111/tpj.17214.

11. Sannier M, Benmamar S, Collemare J, Lemaire C, Caffier V, Duplaix C, et al. Ancient MAX effector variants of a fungal pathogen evade apoplastic immunity in apple. bioRxiv. 2025;12.25.695362. doi: 10.64898/2025.12.25.695362.

12. Langlands-Perry C, Pitarch A, Lapalu N, Cuenin M, Bergez C, Noly A, et al. Quantitative and qualitative plant-pathogen interactions call upon similar pathogenicity genes with a spectrum of effects. Front Plant Sci. 2023;14:1128546. doi: 10.3389/fpls.2023.1128546.

13. Cook DE, Mesarich CH, Thomma BPHJ. Understanding plant immunity as a surveillance system to detect invasion. Annu Rev Phytopathol. 2015;53:541–63. doi: 10.1146/annurev-phyto-080614-120114.

14. Lo Presti L, Lanver D, Schweizer G, Tanaka S, Liang L, Tollot M, et al. Fungal effectors and plant susceptibility. Annu Rev Plant Biol. 2015;66:513–45. doi: 10.1146/annurev-arplant-043014-114623.

15. Rocafort M, Fudal I, Mesarich CH. Apoplastic effector proteins of plant-associated fungi and oomycetes. Curr Opin Plant Biol. 2020;56:9–19. doi: 10.1016/j.pbi.2020.02.004.

16. Deng CH, Plummer KM, Jones DAB, Mesarich CH, Shiller J, Taranto AP, et al. Comparative analysis of the predicted secretomes of Rosaceae scab pathogens *Venturia inaequalis* and *V. pirina* reveals expanded effector families and putative determinants of host range. BMC Genom. 2017;18(1):339. doi: 10.1186/s12864-017-3699-1.

17. Khajuria YP, Akhoon BA, Kaul S, Dhar MK. Secretomic insights into the pathophysiology of *Venturia inaequalis*: The causative agent of scab, a devastating apple tree disease. Pathogens. 2023;12(1):66. doi: 10.3390/pathogens12010066.

18. Le Cam B, Sargent D, Gouzy J, Amselem J, Bellanger M-N, Bouchez O, et al. Population genome sequencing of the scab fungal species *Venturia inaequalis*, *Venturia pirina*, *Venturia aucupariae* and *Venturia asperata*. G3: Genes, Genomes, Genetics. 2019;9(8):2405–14. doi: 10.1534/g3.119.400047.

19. Rocafort M, Bowen JK, Hassing B, Cox MP, McGreal B, de la Rosa S, et al. The *Venturia inaequalis* effector repertoire is dominated by expanded families with predicted structural similarity, but unrelated sequence, to avirulence proteins from other plant-pathogenic fungi. BMC Biol. 2022;20(1):246. doi: 10.1186/s12915-022-01442-9.

20. Thakur K, Chawla V, Bhatti S, Swarnkar MK, Kaur J, Shankar R, et al. *De novo* transcriptome sequencing and analysis for *Venturia inaequalis*, the devastating apple scab pathogen. PLoS One. 2013;8(1):e53937. doi: 10.1371/journal.pone.0053937.

21. Bhagta S, Bhardwaj V, Kant A. Exogenous dsRNA trigger RNAi in *Venturia inaequalis* resulting in down regulation of target genes and growth reduction. Mol Biol Rep. 2023;50(10):8421–9. doi: 10.1007/s11033-023-08736-3.

22. Bowen JK, Mesarich CH, Rees-George J, Cui W, Fitzgerald A, Win J, et al. Candidate effector gene identification in the ascomycete fungal phytopathogen *Venturia inaequalis* by expressed sequence tag analysis. Mol Plant Pathol. 2009;10(3):431–48. doi: 10.1111/j.1364-3703.2009.00543.x.

23. Kucheryava N, Bowen JK, Sutherland PW, Conolly JJ, Mesarich CH, Rikkerink EHA, et al. Two novel *Venturia inaequalis* genes induced upon morphogenetic differentiation during infection and *in vitro* growth on cellophane. Fungal Genet Biol. 2008;45(10):1329–39. doi: 10.1016/j.fgb.2008.07.010.

24. Mesarich CH, Schmitz M, Tremouilhac P, McGillivray DJ, Templeton MD, Dingley AJ. Structure, dynamics and domain organization of the repeat protein Cin1 from the apple scab fungus. Biochim Biophys Acta - Proteins Proteom. 2012;1824(10):1118–28. doi: 10.1016/j.bbapap.2012.06.015.

25. Shiller J, Van de Wouw AP, Taranto AP, Bowen JK, Dubois D, Robinson A, et al. A large family of *AvrLm6*-like genes in the apple and pear scab pathogens, *Venturia inaequalis* and *Venturia pirina*. Front Plant Sci. 2015;6:980. doi: 10.3389/fpls.2015.00980.

26. Patocchi A, Wehrli A, Dubuis P-H, Auwerkerken A, Leida C, Cipriani G, et al. Ten years of VINQUEST: First insight for breeding new apple ,cultivars with durable apple scab resistance. Plant Dis. 2020;104(8):2074–81. doi: 10.1094/pdis-11-19-2473-sr.

27. Lemaire C, De Gracia M, Leroy T, Michalecka M, Lindhard-Pedersen H, Guerin F, et al. Emergence of new virulent populations of apple scab from nonagricultural disease reservoirs. New Phytol. 2016;209(3):1220–9. doi: 10.1111/nph.13658.

28. Chen S, Songkumarn P, Venu RC, Gowda M, Bellizzi M, Hu J, et al. Identification and characterization of *in planta*–expressed secreted effector proteins from *Magnaporthe oryzae* that induce cell death in rice. Mol Plant Microbe Interact. 2013;26(2):191–202. doi: 10.1094/mpmi-05-12-0117-r.

29. Guo X, Zhong D, Xie W, He Y, Zheng Y, Lin Y, et al. Functional identification of novel cell death-inducing effector proteins from *Magnaporthe oryzae*. Rice. 2019;12(1):59. doi: 10.1186/s12284-019-0312-z.

30. Han M, Wang C, Zhu W, Pan Y, Huang L, Nie J. Extracellular perception of multiple novel core effectors from the broad host-range pear anthracnose pathogen *Colletotrichum fructicola* in the nonhost *Nicotiana benthamiana*. Hortic Res. 2024;11(5). doi: 10.1093/hr/uhae078.

31. Hunziker L, Tarallo M, Gough K, Guo M, Hargreaves C, Loo TS, et al. Apoplastic effector candidates of a foliar forest pathogen trigger cell death in host and non-host plants. Sci Rep. 2021;11:19958. doi: 10.1038/s41598-021-99415-5.

32. Kettles GJ, Bayon C, Canning G, Rudd JJ, Kanyuka K. Apoplastic recognition of multiple candidate effectors from the wheat pathogen *Zymoseptoria tritici* in the nonhost plant *Nicotiana benthamiana*. New Phytol. 2017;213(1):338–50. doi: 10.1111/nph.14215.

33. Raffaello T, Asiegbu FO. Small secreted proteins from the necrotrophic conifer pathogen *Heterobasidion annosum* s.l. (HaSSPs) induce cell death in *Nicotiana benthamiana*. Sci Rep. 2017;7(1):8000. doi: 10.1038/s41598-017-08010-0.

34. Gust AA, Felix G. Receptor like proteins associate with SOBIR1-type of adaptors to form bimolecular receptor kinases. Curr Opin Plant Biol. 2014;21:104–11. doi: 10.1016/j.pbi.2014.07.007.

35. Huang WRH, Joosten MHAJ. Immune signaling: receptor-like proteins make the difference. Trends Plant Sci. 2025;30(1):54–68. doi: 10.1016/j.tplants.2024.03.012.

36. Liebrand TWH, van den Berg GCM, Zhang Z, Smit P, Cordewener JHG, America AHP, et al. Receptor-like kinase SOBIR1/EVR interacts with receptor-like proteins in plant immunity against fungal infection. Proc Natl Acad Sci USA. 2013;110(24):10010–5. doi: 10.1073/pnas.1220015110.

37. Liebrand TWH, van den Burg HA, Joosten MHAJ. Two for all: receptor-associated kinases SOBIR1 and BAK1. Trends Plant Sci. 2014;19(2):123–32. doi: 10.1016/j.tplants.2013.10.003.

38. van der Burgh AM, Postma J, Robatzek S, Joosten MHAJ. Kinase activity of SOBIR1 and BAK1 is required for immune signalling. Mol Plant Pathol. 2019;20(3):410–22. doi: 10.1111/mpp.12767.

39. Yasuda S, Okada K, Saijo Y. A look at plant immunity through the window of the multitasking coreceptor BAK1. Curr Opin Plant Biol. 2017;38:10–8. doi: 10.1016/j.pbi.2017.04.007.

40. Kanzaki H, Saitoh H, Takahashi Y, Berberich T, Ito A, Kamoun S, et al. NbLRK1, a lectin-like receptor kinase protein of *Nicotiana benthamiana*, interacts with *Phytophthora infestans* INF1 elicitin and mediates INF1-induced cell death. Planta. 2008;228(6):977–87. doi: 10.1007/s00425-008-0797-y.

41. Kamoun S, van West P, Vleeshouwers VGAA, de Groot KE, Govers F. Resistance of *Nicotiana benthamiana* to *Phytophthora infestans* is mediated by the recognition of the elicitor protein INF1. Plant Cell. 1998;10(9):1413–25. doi: 10.1105/tpc.10.9.1413.

42. Huang WRH, Schol C, Villanueva SL, Heidstra R, Joosten MHAJ. Knocking out *SOBIR1* in *Nicotiana benthamiana* abolishes functionality of transgenic receptor-like protein Cf-4. Plant Physiol. 2020;185(2):290–4. doi: 10.1093/plphys/kiaa047.

43. Zönnchen J, Gantner J, Lapin D, Barthel K, Eschen-Lippold L, Erickson JL, et al. EDS1 complexes are not required for PRR responses and execute TNL-ETI from the nucleus in *Nicotiana benthamiana*. New Phytol. 2022;236(6):2249–64. doi: 10.1111/nph.18511.

44. de la Rosa S, Schol CR, Ramos Peregrina Á, Winter DJ, Hilgers AM, Maeda K, et al. Sequential breakdown of the *Cf-9* leaf mould resistance locus in tomato by *Fulvia fulva*. New Phytol. 2024;243(4):1522–38. doi: 10.1111/nph.19925.

45. Jones DA, Thomas CM, Hammond-Kosack KE, Balint-Kurti PJ, Jones JDG. Isolation of the tomato *Cf-9* gene for resistance to *Cladosporium fulvum* by transposon tagging. Science. 1994;266(5186):789–93. doi: 10.1126/science.7973631.

46. van Kan JA, van den Ackerveken G, de Wit P. Cloning and characterization of cDNA of avirulence gene *avr9* of the fungal pathogen *Cladosporium fulvum*, causal agent of tomato leaf mold. Mol Plant Microbe Interact. 1991;4(1):52–9. doi: 10.1094/mpmi-4-052.

47. Nielsen H. Predicting secretory proteins with SignalP. In: Kihara D, editor. Protein Function Prediction: Methods and Protocols. New York, NY: Springer New York; 2017. p. 59–73.

48. Abramson J, Adler J, Dunger J, Evans R, Green T, Pritzel A, et al. Accurate structure prediction of biomolecular interactions with AlphaFold 3. Nature. 2024;630(8016):493–500. doi: 10.1038/s41586-024-07487-w.

49. Tarallo M, Mesarich CH, McDougal RL, Bradshaw RE. Foliar pine pathogens from different kingdoms share defence-eliciting effector proteins. Mol Plant Pathol. 2025;26(3):e70065. doi: 10.1111/mpp.70065.

50. Qiu C, Zhang H, Liu Z. *Alternaria solani* core effector Aex59 is a new member of the Alt a 1 protein family and is recognized as a PAMP. Int J Biol Macromol. 2024;278:134918. doi: 10.1016/j.ijbiomac.2024.134918.

51. Liu D, Lun Z, Liu N, Yuan G, Wang X, Li S, et al. Identification and characterization of novel candidate effector proteins from *Magnaporthe oryzae*. J Fungi. 2023;9(5):574. doi: 10.3390/jof9050574.

52. Xue C, Park G, Choi W, Zheng L, Dean RA, Xu J-R. Two novel fungal virulence genes specifically expressed in appressoria of the rice blast fungus. Plant Cell. 2002;14(9):2107–19. doi: 10.1105/tpc.003426.

53. Nie J, Yin Z, Li Z, Wu Y, Huang L. A small cysteine-rich protein from two kingdoms of microbes is recognized as a novel pathogen-associated molecular pattern. New Phytol. 2019;222(2):995–1011. doi: 10.1111/nph.15631.

54. Pi L, Yin Z, Duan W, Wang N, Zhang Y, Wang J, et al. A G-type lectin receptor-like kinase regulates the perception of oomycete apoplastic expansin-like proteins. J Integr Plant Biol. 2022;64(1):183–201. doi: 10.1111/jipb.13194.

55. Gong Z, Ning N, Li Z, Xie X, Wilson RA, Liu W. Two *Magnaporthe* appressoria-specific (MAS) proteins, MoMas3 and MoMas5, are required for suppressing host innate immunity and promoting biotrophic growth in rice cells. Mol Plant Pathol. 2022;23(9):1290–302. doi: 10.1111/mpp.13226.

56. Inoue Y, Phuong Vy TT, Singkaravanit-Ogawa S, Zhang R, Yamada K, Ogawa T, et al. Selective deployment of virulence effectors correlates with host specificity in a fungal plant pathogen. New Phytol. 2023;238(4):1578–92. doi: 10.1111/nph.18790.

57. Martínez-Cruz J, Romero D, Hierrezuelo J, Thon M, de Vicente A, Pérez-García A. Effectors with chitinase activity (EWCAs), a family of conserved, secreted fungal chitinases that suppress chitin-triggered immunity. Plant Cell. 2021;33(4):1319–40. doi: 10.1093/plcell/koab011.

58. McCarthy HM, Tarallo M, Mesarich CH, McDougal RL, Bradshaw RE. Targeted gene mutations in the forest pathogen *Dothistroma septosporum* using CRISPR/Cas9. Plants. 2022;11(8):1016. doi: 10.3390/plants11081016.

59. Franco-Orozco B, Berepiki A, Ruiz O, Gamble L, Griffe LL, Wang S, et al. A new proteinaceous pathogen-associated molecular pattern (PAMP) identified in Ascomycete fungi induces cell death in Solanaceae. New Phytol. 2017;214(4):1657–72. doi: 10.1111/nph.14542.

60. Tarallo M, Dobbie KB, Leite LN, Waters TL, Gillard KNT, Sen D, et al. Genomic and culture-based analysis of *Cyclaneusma minus* in New Zealand provides evidence for multiple morphotypes. Phytopathol Res. 2024;6(1):37. doi: 10.1186/s42483-024-00255-8.

61. Tarallo M, McDougal RL, Chen Z, Wang Y, Bradshaw RE, Mesarich CH. Characterization of two conserved cell death elicitor families from the Dothideomycete fungal pathogens *Dothistroma septosporum* and *Fulvia fulva* (syn. Cladosporium fulvum). Front Microbiol. 2022;13:964851. doi: 10.3389/fmicb.2022.964851.

62. Yang Q, Wang J, Sun J, Gao S, Zheng H, Pan Y. A *Fusarium pseudograminearum* secreted protein Fp00392 is a major virulence factor during infection and is recognized as a PAMP1. J Integr Agric. 2025. doi: 10.1016/j.jia.2025.02.030.

63. Nie J, Zhou W, Liu J, Tan N, Zhou J-M, Huang L. A receptor-like protein from *Nicotiana benthamiana* mediates VmE02 PAMP-triggered immunity. New Phytol. 2021;229(4):2260–72. doi: 10.1111/nph.16995.

64. Yang Y, Steidele CE, Rössner C, Löffelhardt B, Kolb D, Leisen T, et al. Convergent evolution of plant pattern recognition receptors sensing cysteine-rich patterns from three microbial kingdoms. Nat Commun. 2023;14(1):3621. doi: 10.1038/s41467-023-39208-8.

65. Liu J, Peng H, Cui J, Huang W, Kong L, Clarke JL, et al. Molecular characterization of a novel effector expansin-like protein from *Heterodera avenae* that induces cell death in *Nicotiana benthamiana*. Sci Rep. 2016;6(1):35677. doi: 10.1038/srep35677.

66. Vieira P, Nemchinov LG. An expansin-Like candidate effector protein from *Pratylenchus penetrans* modulates immune responses in *Nicotiana benthamiana*. Phytopathology. 2020;110(3):684–93. doi: 10.1094/phyto-09-19-0336-r.

67. Ali S, Magne M, Chen S, Côté O, Stare BG, Obradovic N, et al. Analysis of putative apoplastic effectors from the nematode, *Globodera rostochiensis*, and identification of an expansin-like protein that can induce and suppress host defenses. PLoS One. 2015;10(1):e0115042. doi: 10.1371/journal.pone.0115042.

68. Cosgrove DJ. Plant cell wall loosening by expansins. Annu Rev Cell Dev Biol. 2024;40:329–52. doi: 10.1146/annurev-cellbio-111822-115334.

69. Chruszcz M, Chapman MD, Osinski T, Solberg R, Demas M, Porebski PJ, et al. *Alternaria alternata* allergen Alt a 1: A unique β-barrel protein dimer found exclusively in fungi. J Allergy Clin Immunol. 2012;130(1):241–7.e9. doi: 10.1016/j.jaci.2012.03.047.

70. De Vouge MW, Thaker AJ, A. Curran IH, Zhang L, Muradia G, Rode H, et al. Isolation and expression of a cDNA clone encoding an *Alternaria alternata* Alt a 1 subunit. Int Arch Allergy Immunol. 2009;111(4):385–95. doi: 10.1159/000237397.

71. Haueisen J, Möller M, Seybold H, Small C, Wilkens M, Jahneke L, et al. Comparative analyses of compatible and incompatible host-pathogen interactions provide insight into divergent host specialization of closely related pathogens. Mol Plant Microbe Interact. 2025;38(2):235–51. doi: 10.1094/mpmi-10-24-0133-fi.

72. Jeblick T, Leisen T, Steidele CE, Albert I, Müller J, Kaiser S, et al. *Botrytis* hypersensitive response inducing protein 1 triggers noncanonical PTI to induce plant cell death. Plant Physiol. 2022;191(1):125–41. doi: 10.1093/plphys/kiac476.

73. Liu H, Chen S, Gao X, Qian H, Wang Y, Zhang D, et al. The secreted elicitor protein VmHrp1 from *Valsa mali* activates plant immunity through RLP26 to enhance disease resistance. Plant J. 2025;123(3):e70393. doi: 10.1111/tpj.70393.

74. Seifbarghi S, Borhan MH, Wei Y, Ma L, Coutu C, Bekkaoui D, et al. Receptor-like kinases BAK1 and SOBIR1 are required for necrotizing activity of a novel group of *Sclerotinia sclerotiorum* necrosis-inducing effectors. Front Plant Sci. 2020;11:1021. doi: 10.3389/fpls.2020.01021.

75. Wang B, Yang X, Zeng H, Liu H, Zhou T, Tan B, et al. The purification and characterization of a novel hypersensitive-like response-inducing elicitor from *Verticillium dahliae* that induces resistance responses in tobacco. Appl Microbiol Biotechnol. 2012;93(1):191–201. doi: 10.1007/s00253-011-3405-1.

76. Zhang S, Liu L, Li W, Yin M, Hu Q, Chen S, et al. *Alternaria alternata* effector AaAlta1 targets *Cm*WD40 and participates in regulating disease resistance in *Chrysanthemum morifolium*. PLoS Pathog. 2025;21(3):e1012942. doi: 10.1371/journal.ppat.1012942.

77. Zhang Y, Gao Y, Liang Y, Dong Y, Yang X, Qiu D. *Verticillium dahliae* PevD1, an Alt a 1-like protein, targets cotton PR5-like protein and promotes fungal infection. J Exp Bot. 2018;70(2):613–26. doi: 10.1093/jxb/ery351.

78. Zhang Y, Liang Y, Dong Y, Gao Y, Yang X, Yuan J, et al. The *Magnaporthe oryzae* Alt A 1-like protein MoHrip1 binds to the plant plasma membrane. Biochem Biophys Res Commun. 2017;492(1):55–60. doi: 10.1016/j.bbrc.2017.08.039.

79. Gómez-Casado C, Murua-García A, Garrido-Arandia M, González-Melendi P, Sánchez-Monge R, Barber D, et al. Alt a 1 from *Alternaria* interacts with PR5 thaumatin-like proteins. FEBS Lett. 2014;588(9):1501–8. doi: 10.1016/j.febslet.2014.02.044.

80. Gong S, Tang J, Xiao Y, Li T, Zhang Q. The fungal effector AaAlta1 inhibits PATHOGENESIS-RELATED PROTEIN10-2-mediated callose deposition and defense responses in apple. Plant Physiol. 2024;197(1). doi: 10.1093/plphys/kiae599.

81. Karimi-Jashni M, Maeda K, Yazdanpanah F, de Wit PJGM, Iida Y. An integrated omics approach uncovers the novel effector *Ecp20-2* required for full virulence of *Cladosporium fulvum* on tomato. Front Microbiol. 2022;13:919809. doi: 10.3389/fmicb.2022.919809.

82. Nie H-Z, Zhang L, Zhuang H-Q, Shi W-J, Yang X-F, Qiu D-W, et al. The Secreted Protein MoHrip1 Is Necessary for the Virulence of *Magnaporthe oryzae*. Int J Mol Sci. 2019;20(7):1643. doi: 10.3390/ijms20071643.

83. Garrido-Arandia M, Silva-Navas J, Ramírez-Castillejo C, Cubells-Baeza N, Gómez-Casado C, Barber D, et al. Characterisation of a flavonoid ligand of the fungal protein Alt a 1. Sci Rep. 2016;6:33468. doi: 10.1038/srep33468.

84. Liang Y, Cui S, Tang X, Zhang Y, Qiu D, Zeng H, et al. An asparagine-rich protein Nbnrp1 modulate *Verticillium dahliae* protein PevD1-induced cell death and disease resistance in *Nicotiana benthamiana*. Front Plant Sci. 2018;9:303. doi: 10.3389/fpls.2018.00303.

85. Liang Y, Li Z, Zhang Y, Meng F, Qiu D, Zeng H, et al. Nbnrp1 mediates *Verticillium dahliae* effector PevD1-triggered defense responses by regulating sesquiterpenoid phytoalexins biosynthesis pathway in *Nicotiana benthamiana*. Gene. 2021;768:145280. doi: 10.1016/j.gene.2020.145280.

86. Zhang Y, Gao Y, Wang H-L, Kan C, Li Z, Yang X, et al. *Verticillium dahliae* secretory effector PevD1 induces leaf senescence by promoting ORE1-mediated ethylene biosynthesis. Mol Plant. 2021;14(11):1901–17. doi: 10.1016/j.molp.2021.07.014.

87. Zhou R, Zhu T, Han L, Liu M, Xu M, Liu Y, et al. The asparagine-rich protein NRP interacts with the *Verticillium* effector PevD1 and regulates the subcellular localization of cryptochrome 2. J Exp Bot. 2017;68(13):3427–40. doi: 10.1093/jxb/erx192.

88. Fitzgerald A. Investigation of proteins at the apple scab interface [Ph.D.]: University of Auckland; 2004.

89. Pirc K, Albert I, Nürnberger T, Anderluh G. Disruption of plant plasma membrane by Nep1-like proteins in pathogen–plant interactions. New Phytol. 2023;237(3):746–50. doi: 10.1111/nph.18524.

90. Win J. Molecular quest for avirulence factors in *Venturia inaequalis* [Ph.D.]: University of Auckland; 2003.

91. Win J, Greenwood DR, Plummer KM. Characterisation of a protein from *Venturia inaequalis* that induces necrosis in *Malus* carrying the *Vm* resistance gene. Physiol Mol Plant Pathol. 2003;62(4):193–202. doi: 10.1016/S0885-5765(03)00061-4.

92. Rocafort M, Arshed S, Hudson D, Sidhu JS, Bowen JK, Plummer KM, et al. CRISPR-Cas9 gene editing and rapid detection of gene-edited mutants using high-resolution melting in the apple scab fungus, *Venturia inaequalis*. Fungal Biol. 2022;126(1):35–46. doi: 10.1016/j.funbio.2021.10.001.

93. Bradley EL, Ökmen B, Doehlemann G, Henrissat B, Bradshaw RE, Mesarich CH. Secreted glycoside hydrolase proteins as effectors and invasion patterns of plant-associated fungi and oomycetes. Front Plant Sci. 2022;13:853106. doi: 10.3389/fpls.2022.853106.

94. Caffier V, Patocchi A, Expert P, Bellanger M-N, Durel C-E, Hilber-Bodmer M, et al. Virulence characterization of *Venturia inaequalis* reference isolates on the differential set of *Malus* hosts. Plant Dis. 2015;99(3):370–5. doi: 10.1094/pdis-07-14-0708-re.

95. Parisi L, Fouillet V, Schouten H, Groenwold R, Laurens F, Didelot F, et al. Variability of the pathogenicity of *Venturia inaequalis* in Europe. Acta Hortic. 2004;663(1):107–13. doi: 10.17660/ActaHortic.2004.663.13.

96. Stehmann C, Pennycook S, Plummer KM. Molecular identification of a sexual interloper: The pear pathogen, *Venturia pirina*, has sex on apple. Phytopathology. 2001;91(7):633–41. doi: 10.1094/phyto.2001.91.7.633.

97. Hallgren J, Tsirigos KD, Pedersen MD, Almagro Armenteros JJ, Marcatili P, Nielsen H, et al. DeepTMHMM predicts alpha and beta transmembrane proteins using deep neural networks. bioRxiv. 2022:2022.04.08.487609. doi: 10.1101/2022.04.08.487609.

98. Gíslason MH, Nielsen H, Almagro Armenteros JJ, Johansen AR. Prediction of GPI-anchored proteins with pointer neural networks. Curr Res Biotechnol. 2021;3:6–13. doi: 10.1016/j.crbiot.2021.01.001.

99. Weber E, Engler C, Gruetzner R, Werner S, Marillonnet S. A modular cloning system for standardized assembly of multigene constructs. PLoS One. 2011;6(2):e16765. doi: 10.1371/journal.pone.0016765.

100. Engler C, Gruetzner R, Kandzia R, Marillonnet S. Golden Gate shuffling: A one-pot DNA shuffling method based on type IIs restriction enzymes. PLoS One. 2009;4(5):e5553. doi: 10.1371/journal.pone.0005553.

101. van Huijsduijnen RAMH, Cornelissen BJC, van Loon LC, van Boom JH, Tromp M, Bol JF. Virus-induced synthesis of messenger RNAs for precursors of pathogenesis-related proteins in tobacco. EMBO J. 1985;4(9):2167–71. doi: 10.1002/j.1460-2075.1985.tb03911.x.

102. Hopp TP, Prickett KS, Price VL, Libby RT, March CJ, Pat Cerretti D, et al. A short polypeptide marker sequence useful for recombinant protein identification and purification. Bio/Technology. 1988;6(10):1204–10. doi: 10.1038/nbt1088-1204.

103. Taylor RG, Walker DC, Mclnnes RR. *E. coli* host strains significantly affect the quality of small scale plasmid DNA preparations used for sequencing. Nucleic Acids Res. 1993;21(7):1677–8. doi: 10.1093/nar/21.7.1677.

104. Holsters M, Silva B, Van Vliet F, Genetello C, De Block M, Dhaese P, et al. The functional organization of the nopaline *A. tumefaciens* plasmid pTiC58. Plasmid. 1980;3(2):212–30. doi: 10.1016/0147-619X(80)90110-9.

105. Guo Y, Hunziker L, Mesarich CH, Chettri P, Dupont P-Y, Ganley RJ, et al. DsEcp2-1 is a polymorphic effector that restricts growth of *Dothistroma septosporum* in pine. Fungal Genet Biol. 2020;135:103300. doi: 10.1016/j.fgb.2019.103300.

106. Tarallo M, Hayhurst M, Loo TS, Gerth ML, Bradshaw RE. Heterologous expression of secreted proteins from *Phytophthora* in *Pichia pastoris* followed by protein purification by immobilized metal ion affinity (IMAC). In: Gerth ML, Bradshaw RE, editors. Phytophthora: Methods and Protocols. New York, NY: Springer US; 2025. p. 233–48.

107. Rooney HCE, van’t Klooster JW, van der Hoorn RAL, Joosten MHAJ, Jones JDG, de Wit PJGM. *Cladosporium* Avr2 inhibits tomato Rcr3 protease required for *Cf-2*-dependent disease resistance. Science. 2005;308(5729):1783–6. doi: 10.1126/science.1111404.

108. Meng EC, Goddard TD, Pettersen EF, Couch GS, Pearson ZJ, Morris JH, et al. UCSF ChimeraX: Tools for structure building and analysis. Protein Sci. 2023;32(11):e4792. doi: 10.1002/pro.4792.

109. Kearse M, Moir R, Wilson A, Stones-Havas S, Cheung M, Sturrock S, et al. Geneious Basic: An integrated and extendable desktop software platform for the organization and analysis of sequence data. Bioinformatics. 2012;28(12):1647–9. doi: 10.1093/bioinformatics/bts199.

110. Blum M, Andreeva A, Florentino Laise C, Chuguransky Sara R, Grego T, Hobbs E, et al. InterPro: the protein sequence classification resource in 2025. Nucleic Acids Res. 2024;53(D1):D444–D56. doi: 10.1093/nar/gkae1082.

111. Rocafort M, Srivastava V, Bowen JK, Díaz-Moreno SM, Guo Y, Bulone V, et al. Cell wall carbohydrate dynamics during the differentiation of infection structures by the apple scab fungus, *Venturia inaequalis*. Microbiol Spectr. 2023;11(3):e04219–22. doi: 10.1128/spectrum.04219-22.

